# Uridine-sensitized screening identifies genes and metabolic regulators of nucleotide synthesis

**DOI:** 10.1101/2025.03.11.642569

**Authors:** Abigail Strefeler, Zakery N. Baker, Sylvain Chollet, Rachel M. Guerra, Julijana Ivanisevic, Hector Gallart-Ayala, David J. Pagliarini, Alexis A. Jourdain

## Abstract

Nucleotides are essential for nucleic acid synthesis, signaling, and metabolism, and can be synthesized *de novo* or through salvage. Rapidly proliferating cells require large amounts of nucleotides, making nucleotide metabolism a widely exploited target for cancer therapy. However, resistance frequently emerges, highlighting the need for a deeper understanding of nucleotide regulation. Here, we harness uridine salvage and CRISPR-Cas9 screening to reveal regulators of *de novo* pyrimidine synthesis. We identify several factors and report that pyrimidine synthesis can continue in the absence of coenzyme Q (CoQ), the canonical electron acceptor in *de novo* synthesis. We further investigate NUDT5 and report its conserved interaction with PPAT, the rate-limiting enzyme in purine synthesis. We show that in the absence of NUDT5, hyperactive purine synthesis siphons the phosphoribosyl pyrophosphate (PRPP) pool at the expense of pyrimidine synthesis, promoting resistance to chemotherapy. Intriguingly, the interaction between NUDT5 and PPAT appears to be disrupted by PRPP, highlighting intricate allosteric regulation. Our findings reveal a fundamental mechanism for maintaining nucleotide balance and position NUDT5 as a potential biomarker for predicting resistance to chemotherapy.

## Introduction

Pyrimidines and purines are building blocks of life, and their cellular availability relies on two principal pathways: salvage of nucleosides and nucleobases from dietary uptake and nucleic acid turnover, and *de novo* synthesis from precursors such as amino acids and sugars. The latter pathway is especially crucial in rapidly proliferating cells which must meet increased demands for nucleotides to sustain DNA replication and cell growth^1–3^. Pyrimidine *de novo* synthesis involves the sequential action of three key enzymes (Fig. 1A) starting with the multifunctional protein CAD (carbamoyl-phosphate synthetase II, aspartate transcarbamylase, and dihydroorotase). It is followed by dihydroorotate dehydrogenase (DHODH), localized in the mitochondria, whose activity canonically relies on electron transfer to CoQ, participating in the mitochondrial electron transport chain^4–6^. The final enzyme, uridine monophosphate synthase (UMPS), acts in the cytosol to link the pyrimidine ring with PRPP to form uridine monophosphate (UMP), the precursor of all pyrimidine nucleotides. In contrast, *de novo* biosynthesis of purines is initiated directly on the PRPP backbone through a ten-step pathway beginning with the amidophosphoribosyltransferase PPAT (Fig. 1A). Mutations in the genes required for nucleotide biosynthesis lead to rare genetic disorders that, in the case of pyrimidine deficiency, can be treated by oral supplementation with uridine, the main substrate for pyrimidine salvage^7–9^.

**Figure 1.**
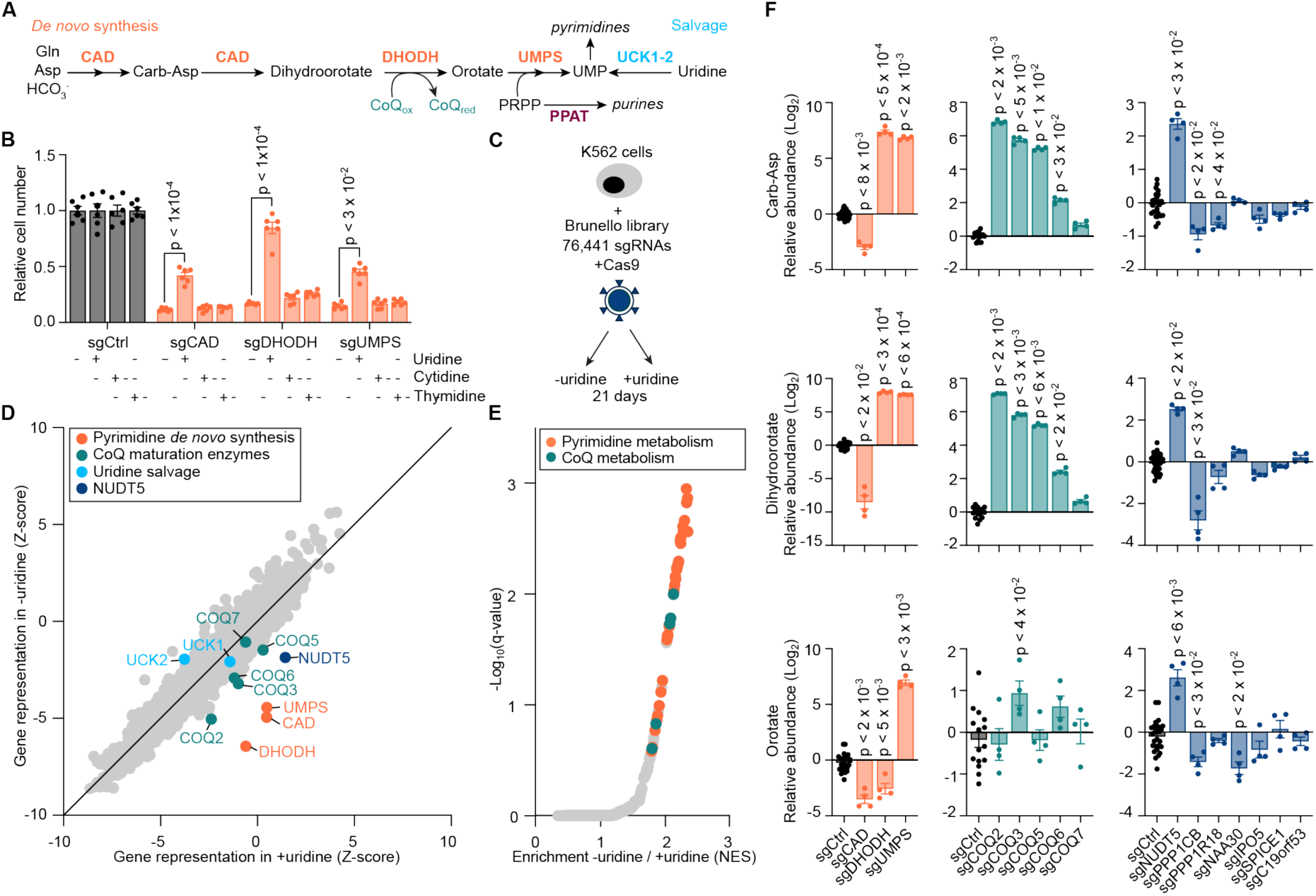
Uridine-sensitized screening identifies players in *de novo* pyrimidine synthesis. **A.** Simplified representation of pyrimidine *de novo* synthesis and salvage pathways. **B.** Proliferation assay of K562 cells transduced with the indicated sgRNAs and Cas9 in medium supplemented individually with 200 µM uridine, 200 µM cytidine, 25 µM thymidine, or left untreated. The lower dose of thymidine was selected due to toxicity. Results are shown as mean ±SEM and p-values are indicated where p < 0.05. Statistical analysis: Student t-test. **C.** Representation of uridine-sensitized genome-wide knockout screen. K562 cells were infected with the Brunello genome-wide sgRNA knockout library, split into medium with or without 200 µM uridine supplementation, and cultured for 21 days. **D.** Gene representation indicated by Z-score in medium with uridine (x-axis) or without uridine (y-axis) from uridine-sensitized knockout screen in K562 cells. **E.** Ranked gene set enrichment analysis using gene ΔZ = Z_-uridine_ – Z_+uridine_ from uridine-sensitized screen and gene ontology biological processes database. The top 50 terms were manually annotated for relationships to pyrimidine or CoQ metabolism. **F.** Relative metabolite abundances in K562 cells transduced with Cas9 and the indicated sgRNAs, normalized to sgCtrl. Results are shown as mean ±SEM and p-values are indicated where p < 0.05. Statistical analysis: non-parametric ANOVA (Kruskall-Wallis test). Abbreviations; Carb-Asp: carbamoyl-aspartate, CoQ_ox_: oxidized coenzyme Q, CoQ_red_: reduced coenzyme Q, NES: normalized enrichment score, PRPP: phosphoribosyl pyrophosphate, sgCtrl: control sgRNA, UMP: uridine monophosphate.

Precise regulation of nucleotide biosynthesis is vital for maintaining nucleotide balance and cellular homeostasis. CAD and PPAT, as the first commitment steps of their respective pathways, are rate-limiting enzymes and are strictly regulated to control *de novo* nucleotide synthesis^4,10–14^. Cancer cells, characterized by rapid proliferation, rely on these pathways to maintain their enhanced metabolic needs, making nucleotide metabolism a prime target for therapeutic intervention. Nucleotide analogs mimic endogenous nucleotides, thereby disrupting DNA replication and RNA stability, and are widely used in cancer and anti-viral therapies^1,15^. However, despite the clinical success of over 20 FDA-approved analogs for treating malignancies such as leukemia and pancreatic cancer, resistance frequently arises due to genetic instability and competition with endogenous substrates^16–18^. These limitations underscore the need to identify novel regulatory mechanisms and therapeutic targets within nucleotide metabolism to overcome resistance and improve treatment outcomes.

Here, we leverage the convergence of pyrimidine *de novo* synthesis and salvage pathways to design a uridine-sensitized CRISPR-Cas9 screening method to identify regulators of pyrimidine *de novo* synthesis. We reveal that mature CoQ is dispensable for pyrimidine *de novo* synthesis and identify the ADP-pyrophosphatase NUDT5 as a pivotal modulator of nucleotide metabolism. We demonstrate that NUDT5 maintains PRPP levels essential for pyrimidine synthesis and nucleobase analog chemotherapies. We further report that NUDT5 acts by inhibiting PPAT through protein-protein interaction in a PRPP-dependent manner, highlighting NUDT5 as a critical and physiological node in ensuring balanced nucleotide production.

## Results

### Uridine-sensitized screening identifies factors in pyrimidine synthesis

To discover genes involved in pyrimidine metabolism, we sought to exploit the dependency on nucleoside salvage exhibited when *de novo* synthesis is impaired^19–21^. We used CRISPR-Cas9 to deplete the three key enzymes required for *de novo* pyrimidine synthesis (*CAD*, *DHODH*, *UMPS*) in K562 myelogenous leukemia cells, which resulted in a strong reduction in proliferation that could be rescued by the addition of uridine to the cell culture medium, but not by addition of downstream nucleosides cytidine nor thymidine (Fig. 1B, Extended Data Fig. 1A). Having confirmed dependency on uridine salvage, we next conducted a genome-wide CRISPR-Cas9 depletion screen comparing cell proliferation in the presence or absence of supplemental uridine (Fig. 1C, Extended Data Fig. 1B, Extended Data Table 1). We applied two analytical methods, a Z-score-based approach^22^ and the Model-based Analysis of Genome-wide CRISPR-Cas9 Knockout (MaGeCK) algorithm^23^, both of which highlighted the three key enzymes of *de novo* pyrimidine biosynthesis as the top essential genes in the absence of uridine, while salvage enzymes (*UCK1, UCK2*) were dispensable (Fig. 1D, Extended Data Fig. 1C, Extended Data Table 1). We confirmed these findings by Gene Set Enrichment Analysis (GSEA)^24,25^ and also found CoQ biosynthesis to be a top-scoring gene ontology class (Fig. 1E, Extended Data Fig. 1D, Extended Data Table 1), an expected result since CoQ is the canonical electron acceptor for DHODH^4–6^ (Fig. 1A). Accordingly, among the known enzymes catalyzing steps in CoQ synthesis, only *COQ7* did not score significantly in our screen (Fig. 1D, Extended Data Fig. 1C). Interestingly, our screen also highlighted other factors not previously linked to pyrimidine biosynthesis, and by prioritizing genes with high scores using both analytical methods we selected *NUDT5*, *PPP1CB*, *PPP1R18*, *NAA30*, *IPO5*, *SPICE1*, and *C19orf53* for further investigation.

Using CRISPR-Cas9, we individually depleted each of these genes, all three *de novo* pyrimidine synthesis enzymes, and five CoQ biosynthetic enzymes, including *COQ7*, in K562 cells. We used targeted metabolomics to analyze the levels of carbamoyl-aspartate, dihydroorotate, and orotate, three intermediates of *de novo* pyrimidine biosynthesis (Fig. 1A), since we reasoned that changes in the abundance of these intermediates would indicate the biosynthetic steps affected in these cells. In validation of this approach, we found altered levels of pyrimidine precursors in cells depleted for each of the three enzymes of *de novo* pyrimidine synthesis, with metabolomes characterized by (I) a profound decrease in all intermediates following *CAD* depletion; (II) accumulation of carbamoyl-aspartate and dihydroorotate, but decreased orotate following *DHODH* depletion; or (III) accumulation of all three intermediates following *UMPS* depletion (Fig. 1F). Analyzing the metabolic profiles of our genes of interest, we found that most fall into one of three major categories: the serine/threonine-protein phosphatase PP1-beta catalytic subunit (*PPP1CB*), its binding partner *PPP1R18*, and to a lesser degree the importin *IPO5*, resembled depletion of *CAD*; *COQ2*, *COQ3*, *COQ5*, *COQ6*, as well as the catalytic subunit of the N-terminal acetyltransferase C (NatC) complex (*NAA30*) resembled depletion of *DHODH*; and the ADP-sugar pyrophosphatase *NUDT5* resembled depletion of *UMPS*, illustrated by the accumulation of all intermediates. *COQ7*, *SPICE1,* and *C19orf53* showed no significant effects (Fig. 1F). Our targeted metabolomics approach validated most of the genes highlighted in our screen and assigned genes to discrete steps in *de novo* pyrimidine synthesis.

### Pyrimidine synthesis in the absence of CoQ

CoQ is the canonical electron acceptor for several enzymes on the inner mitochondrial membrane, including DHODH and the respiratory chain (Fig. 2A). Thus, depletion of CoQ biosynthetic enzymes is expected to block both pyrimidine *de novo* synthesis and respiration^24,26,27^. Our screen effectively revealed and validated most of the central genes in the CoQ biosynthesis pathway, with the notable exception of *COQ7* (Fig. 1F, Extended Data Table 1), which is required for conversion of the CoQ precursor demethoxy-coenzyme Q (DMQ) into demethyl-coenzyme Q (DMeQ), the final intermediate of CoQ biosynthesis^28,29^ (Fig. 2A, B). Given the role of CoQ in DHODH function, the observation that *COQ7* is dispensable for pyrimidine synthesis was unexpected. To investigate this anomaly, we generated K562 single cell knockout clones (*COQ7^KO^*) and measured the levels of DMQ_10_ using lipidomics (Fig. 2C, D). As expected, we found that while DMQ_10_ levels were very low in control cells and in cells depleted for four other *COQ* enzymes, it accumulated strongly in the absence of *COQ7* (Fig. 2D), showing this enzymatic step could not proceed without COQ7. Similarly, we confirmed lower CoQ_10_ levels in all *COQ* knockouts, including *COQ7* depletion (Fig. 2D), and found drastically impaired respiration and failure to thrive in galactose medium, indicating an inability to perform oxidative phosphorylation (OXPHOS) (Fig. 2E, F). This was in marked contrast to the apparent absence of effect of *COQ7* depletion on pyrimidine synthesis, which we quantified using metabolomics and growth assay in uridine-free medium (Fig. 1F, Fig. 2G). Thus, while mature CoQ appears strictly necessary for OXPHOS, we found through our advanced validation of the screen that pyrimidine synthesis is still possible in the absence of its electron acceptor CoQ in *COQ7*-depleted human cells.

**Figure 2.**
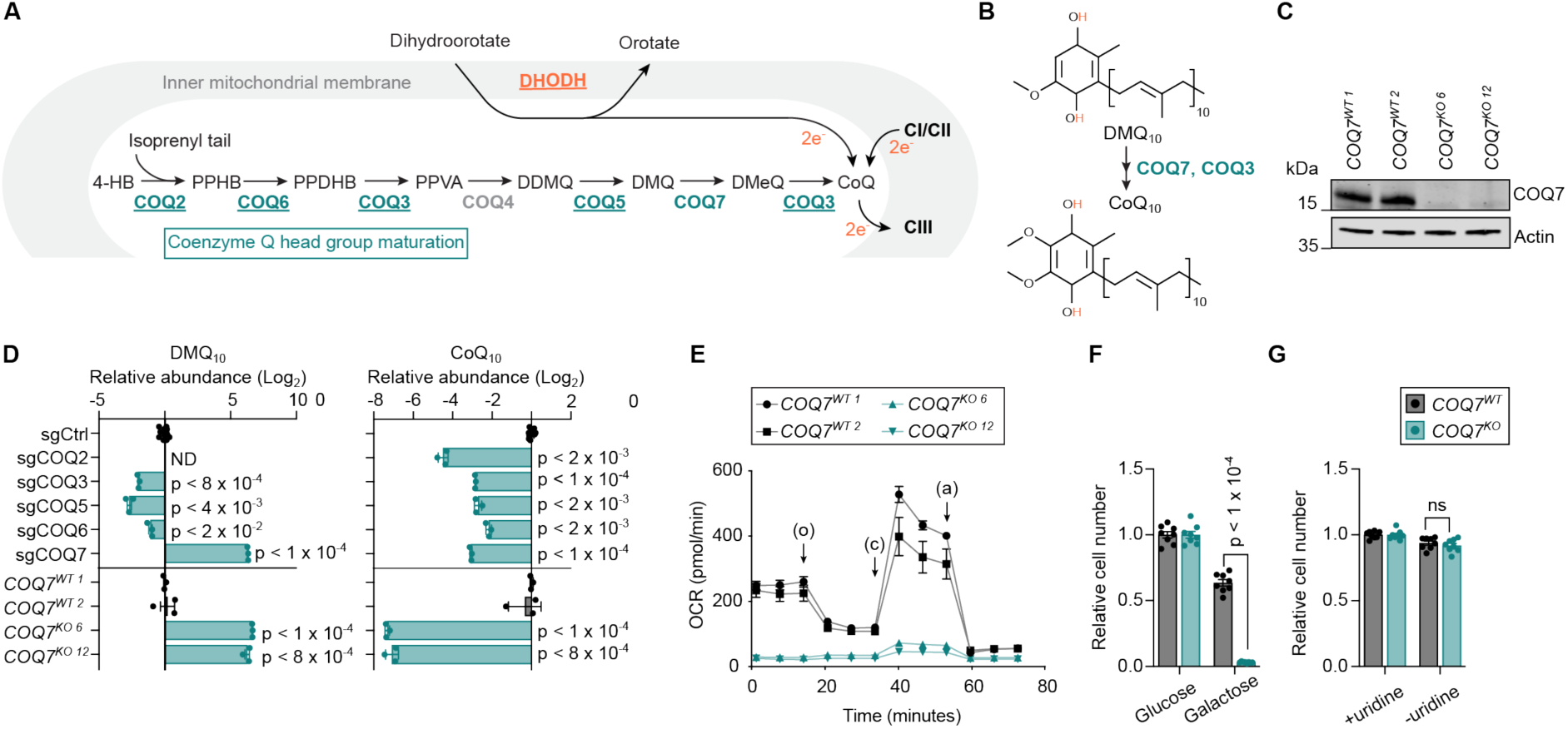
Pyrimidine synthesis, but not OXPHOS, can occur in the absence of *COQ7* and mature CoQ. **A.** Simplified representation of CoQ headgroup maturation reactions within the mitochondria and of mature CoQ as the canonical electron acceptor for DHODH or complexes I/II of the respiratory chain. Enzymes that were present in the library used for screening are colored and further underlined if significant in the screen. **B.** Chemical structures of DMQ_10_ and CoQ_10_. Electron transfer sites are highlighted in orange. **C.** Immunoblot validation of *COQ7^WT^* and *COQ7^KO^* K562 cells. Superscript numbers refer to clone identification. **D.** Relative abundances of DMQ_10_ and CoQ_10_ in K562 cells transduced with the indicated sgRNAs and Cas9 or in single cell clones. Results are normalized to sgCtrl or averaged *COQ7^WT^*samples, respectively. Results are shown as mean ±SEM and p-values are indicated where p < 0.05. Statistical test: one-sample t-test. **E.** Oxygen consumption rate (OCR) on *COQ7* clones. O: oligomycin. C: CCCP. A: antimycin A. **F.** Proliferation assay of *COQ7* clones (two clones each) in glucose- or galactose-supplemented medium. Data are normalized to the respective glucose condition. Results are shown as mean ±SEM and p-value is shown where p < 0.05. Statistical test: Student t-test. **G.** Proliferation assay of *COQ7* clones (two clones each) supplemented with 200 µM uridine or left untreated. Data are normalized to the respective +uridine condition. Results are shown as mean ±SEM. Statistical test: Student t-test. Abbreviations; 4-HB: 4-hydroxybenzoate, CoQ: coenzyme Q, DDMQ: demethoxy-demethyl-coenzyme Q, DMeQ: demethyl-coenyzme Q, DMQ: demethoxy-coenzyme Q, ND: not detected, OCR: oxygen consumption rate, PPDHB: polyprenyl-dihydroxybenzoate, PPHB: polyprenyl-hydroxybenzoate, PPVA: polyprenyl-vanillic acid, sgCtrl: control sgRNA.

### *NUDT5* depletion causes nucleotide imbalance independently of its catalytic activity

The top hit from our screen, apart from the three *de novo* pyrimidine biosynthetic enzymes, was *NUDT5* (*NUDIX5*), encoding a member of the NUDIX (nucleoside diphosphate linked to moiety-X) hydrolase family (Fig. 1D). NUDT5 cleaves ADP-ribose to form ribose 5-phosphate (R5P) and AMP or ATP, depending on phosphate availability, and was also reported to cleave oxidized guanylate nucleotides at high pH^30–32^ (Extended Data Fig. 2A). To gain a broader understanding of the influence of *NUDT5* on cell metabolism, we performed an expanded targeted metabolomics analysis on *NUDT5*-depleted K562 cells and observed significant decrease of mature pyrimidines and accumulation of the intermediates of *de novo* pyrimidine synthesis (Fig. 3A, Extended Data Table 2). In contrast, we found an accumulation of both mature purines and their intermediates, showing nucleotide imbalance (Fig. 3A, B, Extended Data Table 2). We confirmed these findings in *NUDT5*-depleted MCF7 breast cancer cells, a cell line in which earlier work on *NUDT5* was performed^31–33^ (Extended Data Fig. 2B, C).

**Figure 3.**
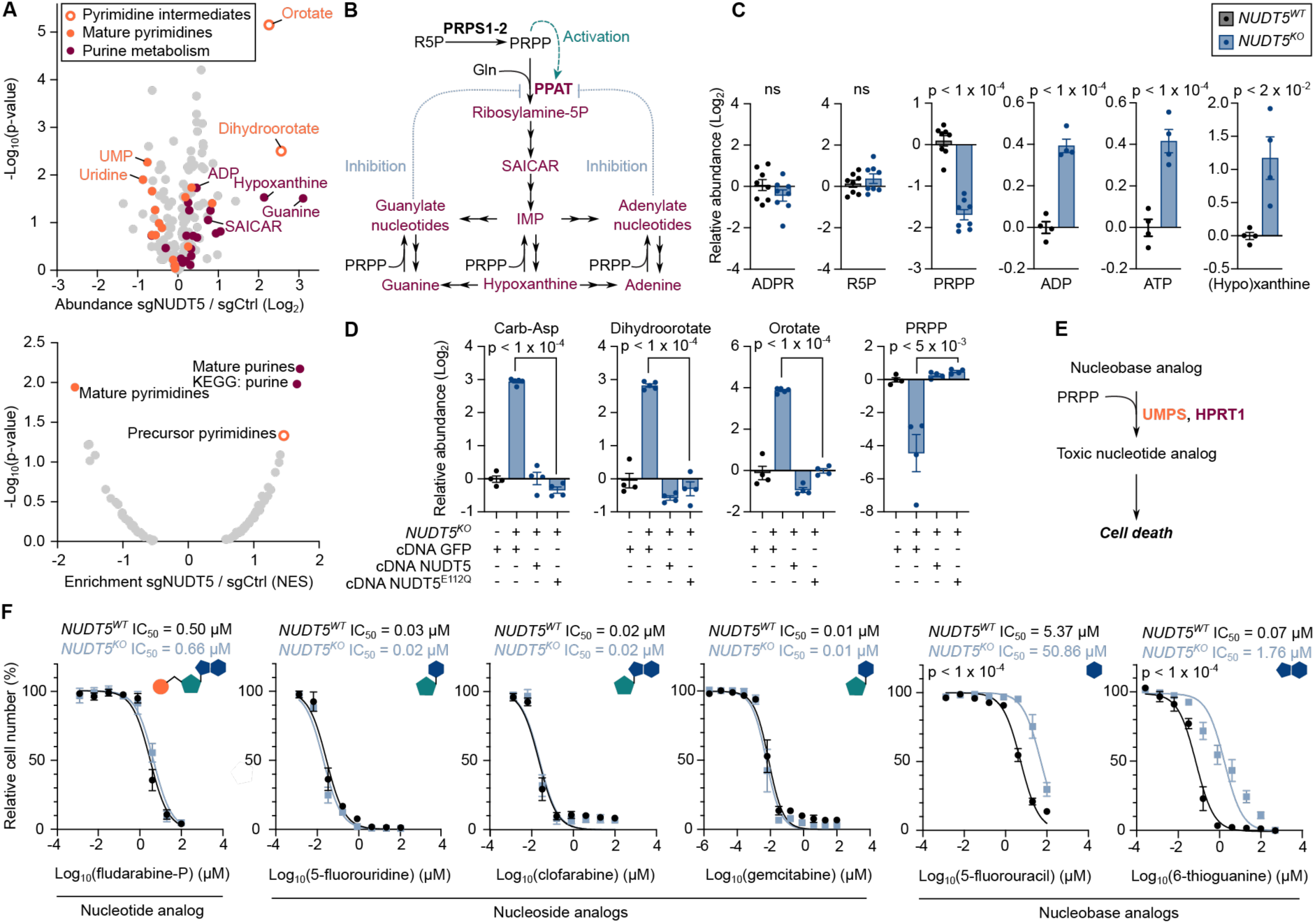
*NUDT5* depletion results in nucleotide imbalance and resistance to nucleobase analog chemotherapy independent of enzymatic activity. **A.** Fold-change in relative metabolite abundances from K562 cells transduced with Cas9 and indicated sgRNAs (top). Enriched metabolic pathways (bottom) were identified by Metabolite Set Enrichment Analysis (MSEA). **B.** Representation of purine biosynthesis including PRPP synthesis. PPAT is regulated by nucleotides (inhibitory feedback, blue dotted lines) and by PRPP (positive feedback, green dashed arrow). **C.** Relative metabolite abundances in *NUDT5* clones (two clones each) as detected using luminescence assays for ADP or ATP, fluorescence assay for (hypo)xanthine, or targeted metabolomics for ADPR, R5P, and PRPP. Data were normalized to *NUDT5^WT^* and results are shown as mean ±SEM with p-values indicated where p < 0.05. Statistical test: Student t-test. **D.** Relative metabolite abundances in *NUDT5* clones complemented with indicated cDNAs. NUDT5^E112Q^ is a catalytic inactive mutant of NUDT5. Data were normalized to *NUDT5^WT^*+GFP cDNA and results are shown as mean ±SEM with p-values indicated where p < 0.05. Statistical test: Student t-test. **E.** Representation of nucleobase analog conversion into toxic compounds using PRPP. **F.** Proliferation assay of *NUDT5* clones (two clones each) in response to increasing dose of purine and pyrimidine nucleotide analogs. A molecular representation is shown with the nitrogenous base (blue polygon), ribose sugar (green pentagon), and phosphate (orange circle). Results are shown as mean ±SEM and are fitted to a non-linear regression using the least-squares method. IC_50_ values are indicated for all curves and p-values are indicated where adjusted p < 0.05. Statistical test: extra sum-of-squares F-test with Bonferroni correction. Abbreviations; ADPR: ADP-ribose, Carb-Asp: carbamoyl-aspartate, IC_50_: half-inhibitory concentration, IMP: inosine monophosphate, NES: normalized enrichment score, PRPP: phosphoribosyl pyrophosphate, R5P: ribose-5-phosphate, SAICAR: 5’-phosphoribosyl-4-(N-succinylcarboxamide)-5-aminoimidazole, sgCtrl: control sgRNA, UMP: uridine monophosphate.

To extend our investigation, we generated *NUDT5^KO^* K562 single cell clones (Extended Data Fig. 2D), in which we confirmed altered pyrimidine and purine metabolism using both targeted metabolomics and orthogonal biochemical assays (Fig. 3C, Extended Data Fig. 2E). In addition, we measured ADP-ribose and R5P, two main metabolites of the NUDT5 reaction, but found no differences in their abundance, indicating that ADP-ribose catabolism from NUDT5 does not participate significantly to these pools (Fig. 3C). Having observed a *UMPS*-like phenotype for *NUDT5* (Fig. 1F), we next analyzed the ribose donor PRPP and found that its levels were significantly decreased in *NUDT5^KO^* cells (Fig. 3C). This observation explains the decreased pyrimidine synthesis seen upon *NUDT5* depletion, but was in part unexpected, since PRPP is also the precursor to purine synthesis (Fig. 3B).

We next tested whether the catalytic activity of NUDT5 was required for its function in nucleotide metabolism. To address this question, we expressed a wild-type or a catalytically inactive mutant (E112Q)^32,34^ of NUDT5 in *NUDT5^KO^* cells (Extended Data Fig. 2F). We found that the levels of PRPP and pyrimidine synthesis intermediates returned to baseline in the presence of either wild-type or mutant NUDT5 (Fig. 3D), suggesting that NUDT5 catalytic activity is not essential for its effect on pyrimidine synthesis. We confirmed these findings using an orthogonal approach based on the inhibition of endogenous NUDT5 with the nanomolar inhibitor TH5427^35^, in which we also observed no effect on nucleotide metabolism (Extended Data Fig. 2G, H). Together, our results indicate that whereas PRPP and mature pyrimidine pools are low in *NUDT5*-depleted cells, purine synthesis appears to function at a higher rate, suggesting preferential mobilization of the PRPP pool towards purines, with a detrimental effect on pyrimidine synthesis. Curiously, the role of NUDT5 in maintaining nucleotide balance appears to be independent of its catalytic activity.

### Loss of *NUDT5* protects against nucleobase analog chemotherapy

Having seen nucleotide imbalance and a strong effect on the PRPP pool in *NUDT5^KO^* cells, we next tested whether *NUDT5* could affect the sensitivity of cancer cells to nucleobase analog drugs, which require PRPP for their conversion into toxic nucleotide analogs (Fig. 3E). This hypothesis is consistent with the results of two high-throughput screens that identified a role for *NUDT5* in 6-thioguanine resistance^36,37^, although in both cases the mechanism of resistance remained unclear. We first determined the half-maximal inhibitory concentration (IC_50_) of 5-fluorouracil (5-FU), a pyrimidine nucleobase analog partly metabolized by UMPS^38^ and approved for cancer therapy, and observed that *NUDT5^KO^* cells were an order of magnitude more resistant to 5-FU than their corresponding wild-type counterparts (Fig. 3F). In contrast, sensitivity to 5-fluorouridine, the nucleosidic form of 5-FU that does not require PRPP for processing, was unchanged. We extended these observations to four other approved purine and pyrimidine analogs and again observed that while *NUDT5^KO^* cells remained equally sensitive to all tested nucleotide and nucleoside analogs, they were consistently more resistant to nucleobase analogs that rely on PRPP, irrespective of whether the molecules were purine or pyrimidine based. Our results indicate that *NUDT5* depletion promotes resistance to nucleobase analog therapies and positions NUDT5 as a potential biomarker for therapy efficacy.

### NUDT5 restrains PPAT to support pyrimidine synthesis

Having excluded an enzymatic mechanism through which NUDT5 affects pyrimidine synthesis, we investigated changes in the proteome that occur upon *NUDT5* depletion. However, no substantial changes were observed in the abundance of enzymes involved in purine, pyrimidine, or PRPP synthesis (Extended Data Fig. 3A, Extended Data Table 3). We then immunoprecipitated FLAG-tagged NUDT5 and investigated its binding partners by mass spectrometry (Fig. 4A, Extended Data Table 3). We found only few NUDT5-interacting proteins, notably among them PPAT, the rate-limiting and first committed enzyme required for *de novo* purine biosynthesis^12^ (Fig. 3B). An interaction between NUDT5 and PPAT has previously been reported in proteome-scale protein-protein interaction studies^39–41^ and we used immunoprecipitation with either of NUDT5-FLAG wild-type, its catalytic inactive E112Q mutant, or PPAT-FLAG as bait to confirm the association between NUTD5 and PPAT (Fig. 4B, Extended Data Fig. 3B).

**Figure 4.**
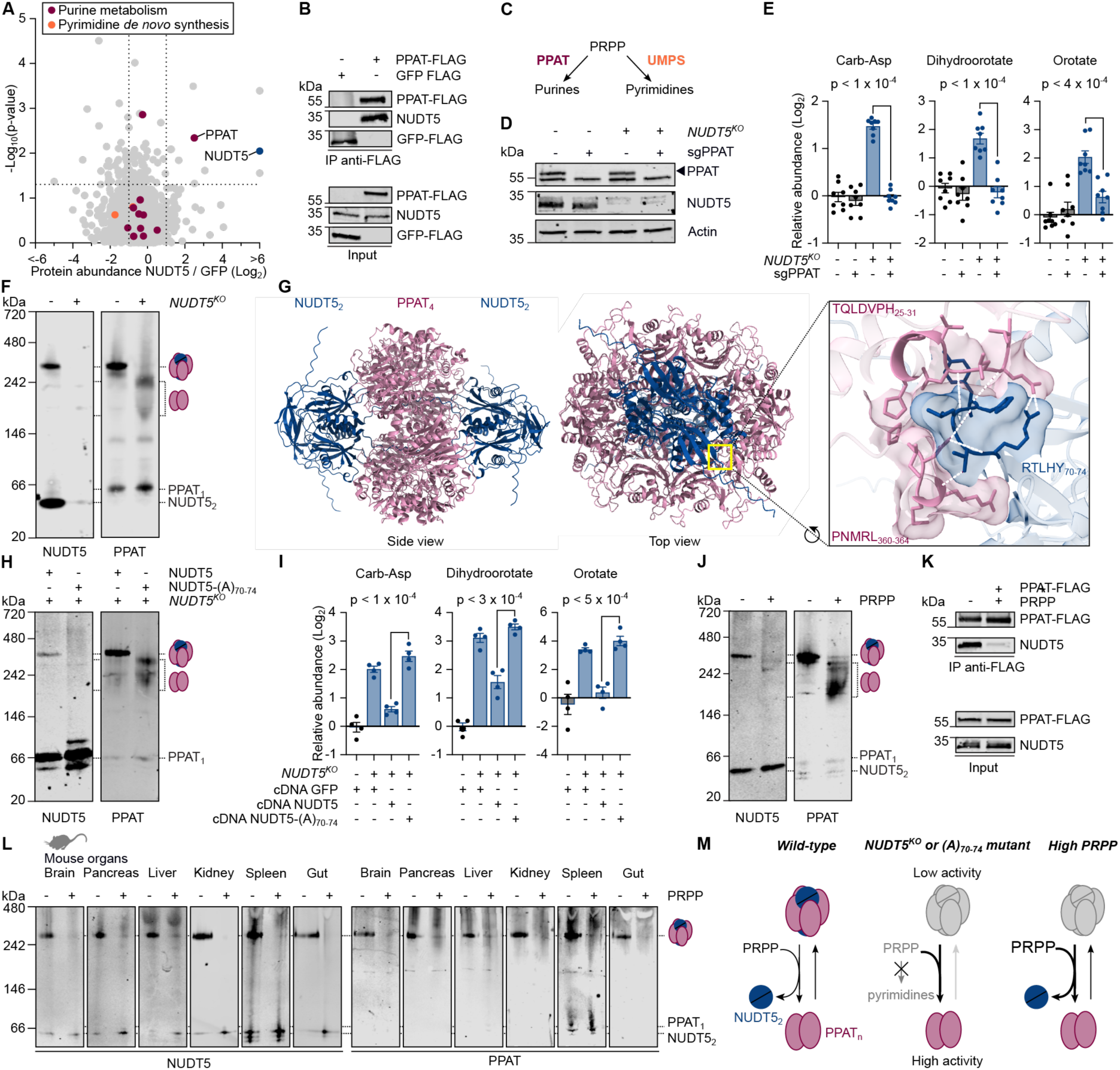
NUDT5 interacts with PPAT and prevents excessive PPAT-mediated purine synthesis at the expense of pyrimidines. **A.** Endogenous protein binding partners from immunoprecipitation of FLAG-tagged NUDT5 in 293T cells followed by mass spectrometry-based proteomics. Statistical test: Student t-test. **B.** Co-immunoprecipitation from 293T cells using FLAG-tag as bait. **C.** Schematic representation of PRPP use for purine or pyrimidine *de novo* synthesis. **D.** Representative immunoblot of *NUDT5* clones transduced with Cas9 and sgRNA against PPAT or control. **E.** Relative metabolite abundances in *NUDT5* clones (two clones each) transduced with Cas9 and sgRNA against PPAT or control. Data were normalized to *NUDT5^WT^* sgCtrl and results are shown as mean ±SEM with p-values indicated where p < 0.05. Statistical test: Student t-test. **F.** Parallel native PAGE on *NUDT5* clones. **G.** AlphaFold 3 prediction of a complex consisting of PPAT tetramer (pink) and two NUDT5 dimers (blue), showing a zoom to the interaction interface of one NUDT5 and two PPAT molecules with amino acid residues and hydrogen bonds (dashed white lines) indicated. **H.** Parallel native PAGE on *NUDT5* clones with indicated cDNA complementation. **I.** Relative metabolite abundances in *NUDT5* clones with indicated cDNA complementation. Data are normalized to *NUDT5^WT^* +GFP cDNA and results are shown as mean ±SEM with p-values indicated where p < 0.05. Statistical test: Student t-test. **J.** Parallel native PAGE on K562 cells with 10 mM PRPP in the lysis buffer or left untreated. **K.** Co-immunoprecipitation from 293T cells using FLAG-tagged PPAT as bait with 10 mM PRPP in the buffer or left untreated. **L.** Parallel native PAGE on mouse organs with 10 mM PRPP in the lysis buffer or left untreated. **M.** Proposed model of PPAT regulation. NUDT5 binding promotes formation of the large low-activity PPAT form. When interaction with NUDT5 is disrupted, PPAT constitutively forms the small high-activity form, depleting PRPP and hindering pyrimidine synthesis. In physiological conditions, elevated PRPP similarly promotes complex dissociation and PPAT activity for purine synthesis. Abbreviations; Carb-Asp: carbamoyl-aspartate, PRPP: phosphoribosyl pyrophosphate.

PPAT and UMPS both consume and potentially compete for PRPP, a metabolite we found to be critically low in *NUDT5*-depleted cells (Fig. 3C, 4C). To determine whether PPAT may be at the origin of the pyrimidine deficiency seen in these cells we sought to simultaneously deplete *PPAT* and *NUDT5* in an epistasis genetic experiment. However, similar to uridine dependency when pyrimidine *de novo* synthesis is impossible (Fig. 1B), *PPAT*-depleted cells become dependent on purine salvage, and we therefore supplemented these with inosine (Extended Data Fig. 3C). Importantly, we found that *PPAT* depletion was sufficient to restore carbamoyl-aspartate, dihydroorotate, and orotate to wild-type levels in *NUDT5^KO^* cells (Fig. 4D, E). Our observations indicate a contribution for PPAT to the pyrimidine phenotype and position PPAT downstream of NUDT5.

Seminal work from the 1970s identified the presence of PPAT in two interconvertible forms, consisting in a larger and partially inactive ∼300 kDa form, and a smaller active form. We thus further investigated the NUDT5-PPAT interaction using native gel electrophoresis in K562 cells, where we majoritarily observed the large ∼300 kDa form with low enzymatic activity^42–44^ (Fig. 4F). Interestingly, this large PPAT oligomer co-migrated with NUDT5, which was also present independently as dimer^32^ (Fig. 4F). Strikingly however, when *NUDT5* was depleted, the PPAT complex dissociated (Fig. 4F), forming smaller oligomers consistent with the active form^42–44^, and suggesting the interaction is inhibitory for PPAT, as hinted to by our genetic experiment (Fig. 4D, E). We next aimed to model the NUDT5-PPAT complex, but no experimental structure of human PPAT currently exists, likely due to its labile iron-sulfur cluster and sensitivity to oxygen that have hindered its recombinant production^45^. We thus used AlphaFold 3^46^ to predict the structure of a human PPAT monomer (57 kDa) and tetramer (228 kDa), both of which exhibited strong similarities to experimentally-determined bacterial PPAT homolog purF^47,48^ (Extended Data Figure 3D). We next added one or two NUDT5 dimers and found that these could cap either side of the PPAT barrel (Fig. 4G, Extended Data Fig. 3E, Extended Data Movie 1). The predicted interaction interfaces consisted of NUDT5 residues RTLHY_70-74_ that extended between two PPAT molecules, associated with residues TQLDVPH_25-31_ on one, and PNMRL_360-364_ on the other (Fig. 4G), all of which are conserved across vertebrates (Extended Data Fig. 3F). To determine the validity of the proposed model, we next mutated NUDT5 residues 70-74 to alanines and reintroduced the corresponding cDNA into *NUDT5^KO^* cells (Extended Data Fig. 3G). Importantly, we found that NUDT5-(A)_70-74_ was unable to rescue the formation of the NUDT5-PPAT complex nor levels of pyrimidine precursors (Fig. 4H, I), consistent with the AlphaFold predictions (Extended Data Fig. 3E), and showing that NUDT5 must physically interact with PPAT to support pyrimidine synthesis.

Finally, we investigated whether the NUDT5-PPAT interaction could be itself regulated. In addition to its role as a substrate, PRPP is an established allosteric activator of PPAT that influences its oligomerization, as addition of PRPP to cell lysates is sufficient to convert PPAT’s large form in its active smaller form^42–44,49^. Strikingly, and consistent with these reports, we found that PRPP could induce formation of smaller PPAT oligomers that were similar to those seen in *NUDT5^KO^* cells (Fig. 4F, J). Accordingly, we found that PRPP also disrupted the interaction between both proteins across ten cell lines and six mouse organs, indicating how a physiological activator can displace NUDT5 from PPAT (Fig. 4J-M, Extended Data Fig. 3H). Together, our genetic and biochemical observations demonstrate that NUDT5 interacts with PPAT to promote the formation of a high-molecular weight, low-activity PPAT form that can be disrupted by the allosteric activator PRPP, and that in *NUDT5*-depleted cells, constitutively active low-molecular weight form PPAT consumes the endogenous PRPP pool to synthesize purines, at the expense of *de novo* pyrimidine synthesis (Fig. 4M).

## Discussion

By combining uridine sensitization and genome-wide CRISPR-Cas9 screening, we discovered factors required for *de novo* pyrimidine synthesis. While our approach successfully recovered *CAD, DHODH,* and *UMPS* as the most important factors, it also highlighted several other genes that we extensively validated using metabolomics and lipidomics, assigning genes to specific steps in pyrimidine synthesis.

Our findings challenge the presumed indispensability of CoQ for pyrimidine synthesis. We showed that pyrimidine synthesis persists in *COQ7*-deficient cells, despite marked CoQ depletion. The accumulation of DMQ in these cells and its molecular resemblance to CoQ (Fig. 2) suggest that DMQ may substitute as an electron acceptor for DHODH. Recent research has highlighted rhodoquinone as an alternative electron carrier able to maintain pyrimidine synthesis in the absence of CoQ^50^, and DMQ may play a similar role, as it is interesting to note that CoQ, DMQ, and rhodoquinone differ from a single chemical group on the same position of the polar head group. In contrast however, while rhodoquinone can also maintain OXPHOS, we found that *COQ7* loss severely impairs respiration (Fig. 2), indicating limited ability of DMQ to function as an electron carrier for mitochondrial complexes. While additional work, notably using chemically synthesized DMQ, will be required to validate electron transfer from DHODH to DMQ, our results hint at structural differences in CoQ-binding sites and may pave the way for increasingly specific DHODH inhibitors.

Through genetic screening and metabolomics, we identified NUDT5 as a mediator of nucleotide balance. We showed that NUDT5 is an inhibitory binding partner to the rate-limiting purine enzyme PPAT, and that its loss promotes purine biosynthesis, depleting the endogenous PRPP pool, and thus impairing pyrimidine *de novo* synthesis and nucleobase analog metabolism. Intriguingly, while PRPP also acts as an allosteric activator for PPAT^42–44,49^, our findings expand the view of PPAT regulation by identifying NUDT5 and showing that its binding is sensitive to PRPP treatment. Further research, possibly including detailed structural analysis, will be needed to clarify the interplay between NUDT5- and PRPP-mediated PPAT regulation and investigate their full impact on nucleotide synthesis and macromolecular structures such as the purinosome^51^. Furthermore, nutrient availability, already known to affect PRPP levels and nucleotide synthesis^13,14,52,53^, may represent a driving factor behind this fundamental regulatory mechanism and will merit further investigation.

Our findings also positioned NUDT5 in the context of anti-cancer treatments and suggest a future role as a biomarker for efficacy of nucleobase analog therapy. In addition, NUDT5 expression has previously been linked with cancer proliferation and malignancy^32,33,35,54,55^, and while its reported role in DNA damage repair^32,33,35^ should be considered, our findings on the role of NUDT5 as a safeguard of nucleotide balance suggest a dual protective function against genetic instability, indicating a promising therapeutic target. Curiously however, we found that the known catalytic function of NUDT5 was not implicated in its role in nucleotide synthesis, and further characterization of both enzymatic and structural roles and their respective impact on tumorigenesis will be a focus for further research.

Our work highlights the power of nucleoside-sensitized genetic screens to identify genes involved in nucleotide metabolism and human disease. We expect our approach can be extended to investigate other pathways of nucleotide metabolism, for example by screening in inosine-containing conditions (Extended Data Fig. 3C). Together, our findings are highly relevant to understanding the limitations, and improving the effectiveness, of nucleotide analog-based cancer therapies.

## Acknowledgements

The authors would like to thank the past and present members of the Jourdain laboratory, Roni Wright, the Metabolomics Platform, and the Protein Analysis Facility (University of Lausanne). This work was supported by a grant from the Swiss National Science Foundation (310030_200796 to AAJ), an NIH award (R35GM131795 to D.J.P.), funds from the BJC Investigator Program (to D.J.P.), and a grant from the Emma Muschamp Foundation (to AAJ). D.J.P. is an investigator of the Howard Hughes Medical Institute.

## Extended Data and Legends

**Extended Data Fig. 1.**
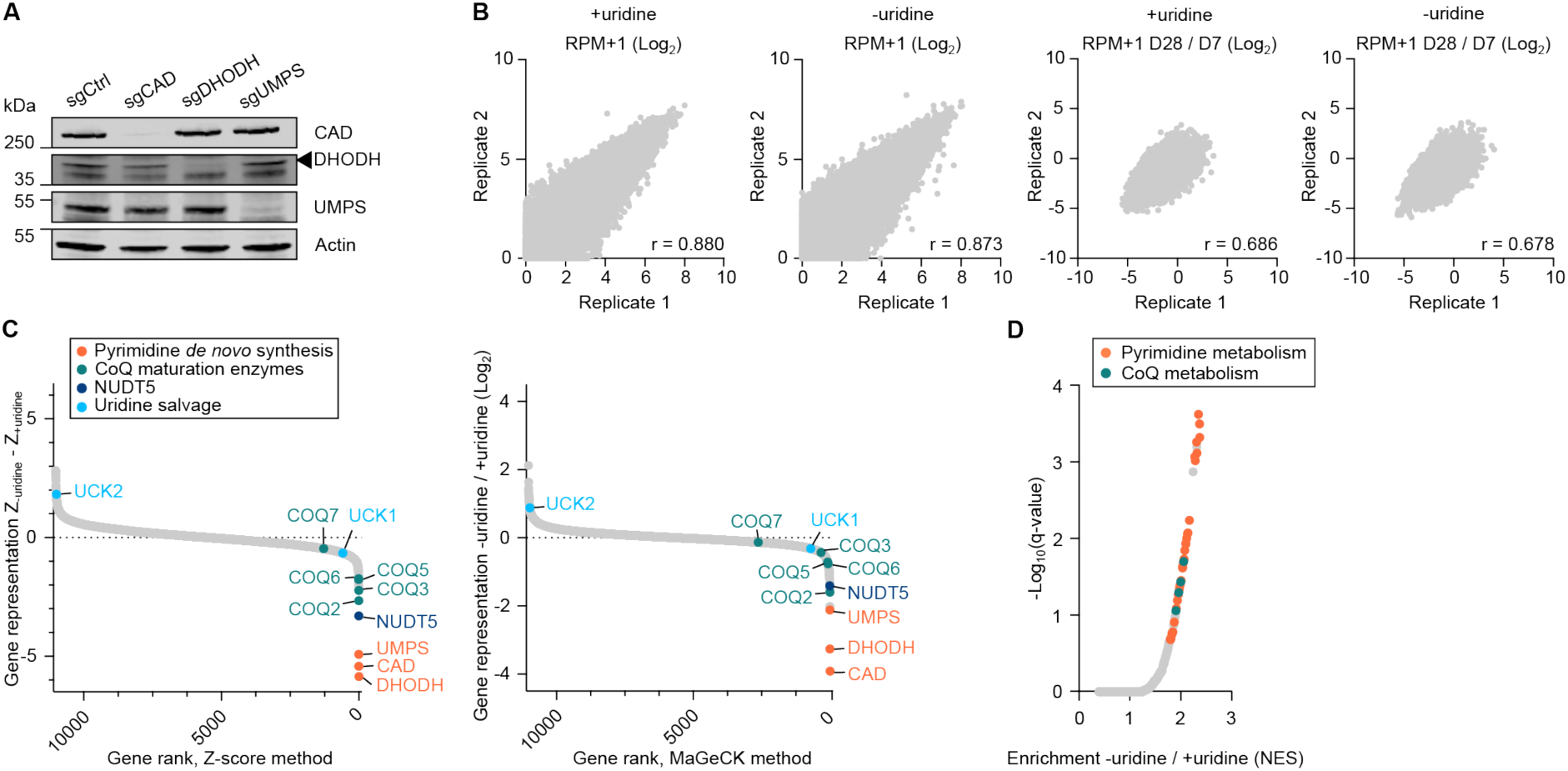
Uridine-sensitized screening discovers genes in pyrimidine synthesis. **A.** Immunoblot validation for knockout of the three key enzymes of pyrimidine *de novo* synthesis in K562 cells. **B.** Comparison of sgRNA representation in replicate infections of CRISPR-Cas9 screening. Data are represented as RPM+1 or as a fold-change with day 7 post-infection (day of media switch). Each point is one sgRNA of an expressed gene. Statistical test: Pearson’s correlation. **C.** Gene representation in medium without uridine supplementation relative to medium with 200 µM uridine, as calculated by Z-score (left) or MaGeCK (right) analysis of uridine-sensitized screen. **D.** Ranked gene set enrichment analysis using gene fold-changes from MaGeCK analysis of uridine-sensitized screen and gene ontology biological processes database. Top 50 terms were manually annotated for relationships to pyrimidine and CoQ metabolism. Abbreviations; CoQ: coenzyme Q, NES: normalized enrichment score, RPM: reads per million, sgCtrl: control sgRNA.

**Extended Data Fig. 2.**
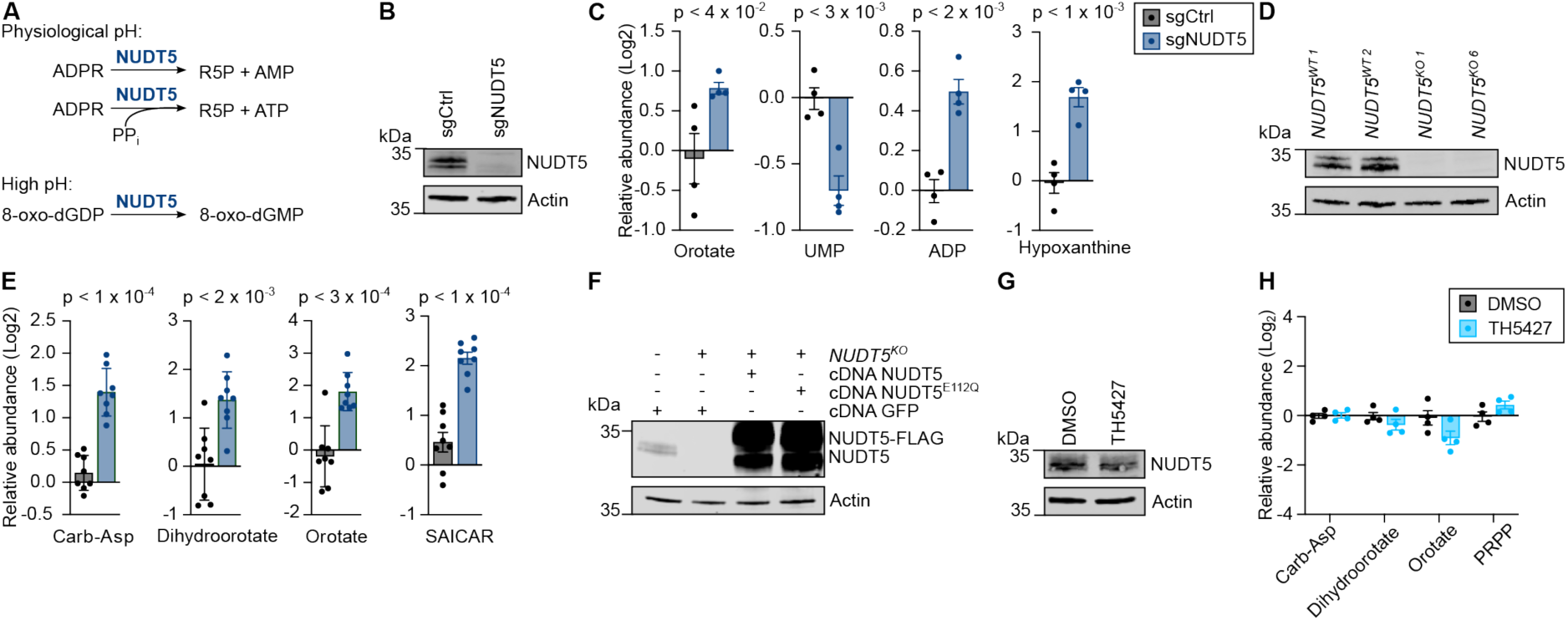
*NUDT5* depletion induces nucleotide imbalance. **A.** Representation of reported NUDT5 enzymatic activities. **B.** Immunoblot validation of NUDT5 knockout in MCF7 cells. **C.** Relative metabolite abundances in MCF7 cells transduced with Cas9 and sgRNAs against NUDT5 or control. Data were normalized to sgCtrl and results are shown as mean ±SEM with p-values indicated where p < 0.05. Statistical test: Student t-test. **D.** Immunoblot validation of *NUDT5^WT^* and *NUDT5^KO^* K562 single cell clones. Superscript numbers refer to clone identifiers. **E.** Relative metabolite abundances in *NUDT5* clones (two clones each). Data were normalized to *NUDT5^WT^* and results are shown as mean ±SEM with p-values indicated where p < 0.05. Statistical test: Student t-test. **F.** Immunoblot validation of indicated cDNA complementation in *NUDT5* clones. **G.** Immunoblot validation of maintained NUDT5 expression following treatment with 10 µM TH5427 or DMSO for 36 h. **H.** Relative metabolite abundances in K562 cells treated with 10 µM TH5427 or DMSO for 36 h. Data were normalized to DMSO-treated and results are shown as mean ±SEM. Abbreviations; 8-oxo-dGDP: oxidized deoxyguanosine diphosphate, ADPR: adenosine diphosphate-ribose, Carb-Asp: carbamoyl-aspartate, PRPP: phosphoribosyl pyrophosphate, R5P: ribose-5-phosphate, SAICAR: 5’-phosphoribosyl-4-(N-succinylcarboxamide)-5-aminoimidazole, sgCtrl: control sgRNA, UMP: uridine monophosphate.

**Extended Data Fig. 3.**
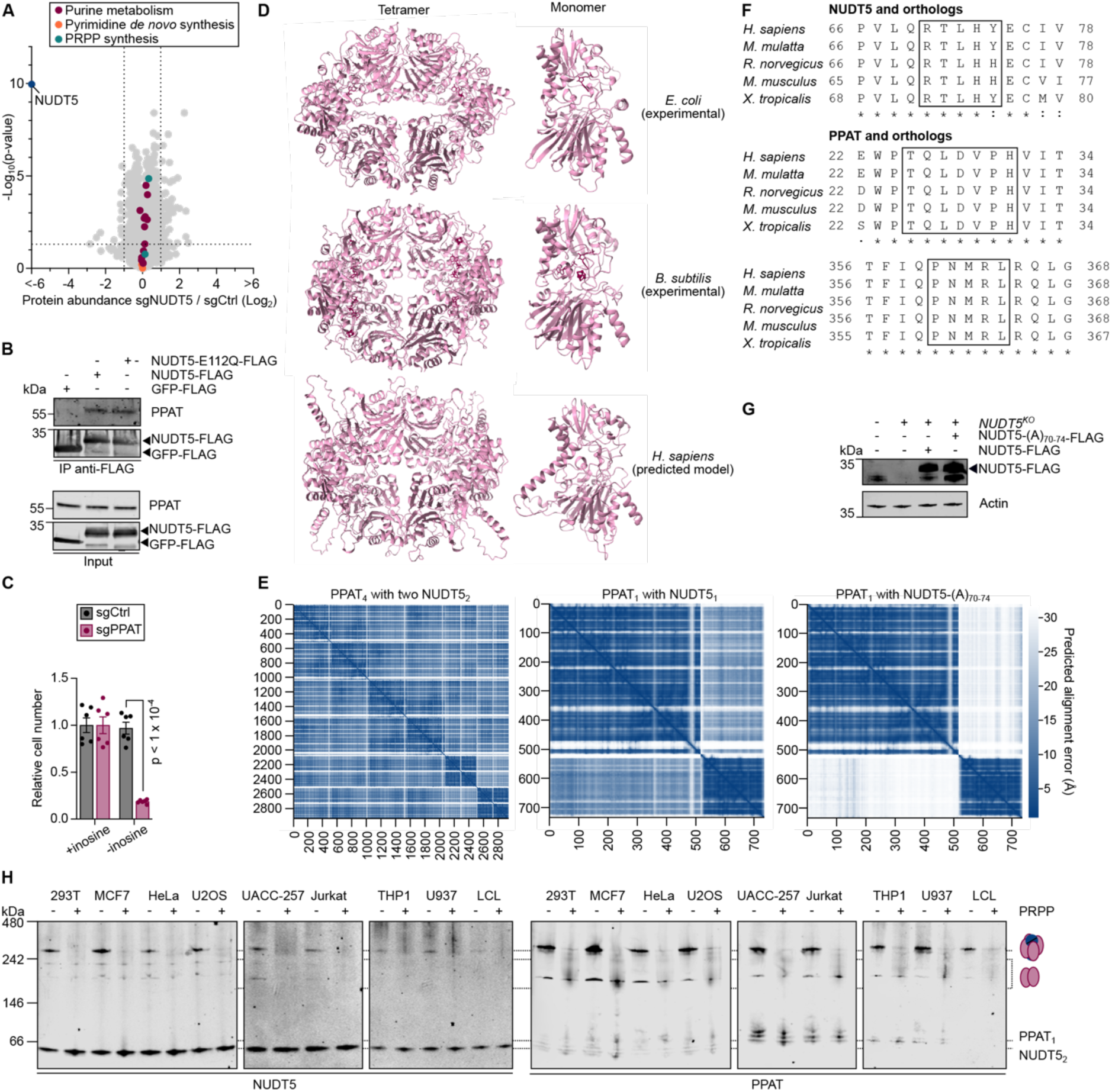
A PPAT-NUDT5 complex is conserved across species and cell types. **A.** Global proteomics comparing K562 cells transduced with Cas9 and sgRNAs against NUDT5 or control. Statistical test: Student t-test. **B.** Co-immunoprecipitation of endogenous PPAT from 293T using FLAG-tagged NUDT5 (wild-type or E112Q mutant) as bait. **C.** Proliferation assay of *NUDT5^WT^* K562 cells transduced with Cas9 and sgRNAs against PPAT or control and grown in medium supplemented with 200 µM inosine or left untreated. Data were normalized to +inosine condition and results are shown as mean ±SEM with p-values indicated where p < 0.05. Statistical test: Student t-test. **D.** Structures of bacterial purF determined experimentally in complex with AMP^47,48^ and AlphaFold 3 predicted structure of human PPAT. Monomers were isolated from the tetramer structures. Abbreviations; PRPP: phosphoribosyl pyrophosphate, sgCtrl: control sgRNA. **E.** AlphaFold 3 predicted alignment error of indicated NUDT5-PPAT complexes. **F.** Sequence alignment of NUDT5 and PPAT proteins across vertebrate species in regions predicted by AlphaFold 3 to mediate the NUDT5-PPAT interaction (black box). **G.** Immunoblot validation of indicated cDNA complementation in *NUDT5* clones. **H.** Parallel native PAGE across seven cell lines treated with 10 mM PRPP in the lysis buffer or left untreated. Abbreviations; PRPP: phosphoribosyl pyrophosphate, sgCtrl: control sgRNA.

**Extended Data Table 1** Data associated with uridine-sensitized CRISPR-Cas9 screen. Data include read count data, Z-score and MaGeCK analyses, and associated Gene Set Enrichment analyses.

**Extended Data Table 2** Data associated with multiple-pathways metabolomics analysis and associated Metabolite Set Enrichment Analysis.

**Extended Data Table 3** Proteomics data including global proteomics on *NUDT5*-depleted cells and IP-MS proteomics for NUDT5-interacting partners compared with GFP control.

**Extended Data Table 4** sgRNA and cDNA sequences used in this study for individual gene depletion or expression.

**Extended Data Table 5** Mass table for targeted metabolomics analysis.

**Extended Data Movie 1** AlphaFold 3 molecular model prediction of PPAT tetrameric complex and association with NUDT5.

**Source Data Fig. 2:**
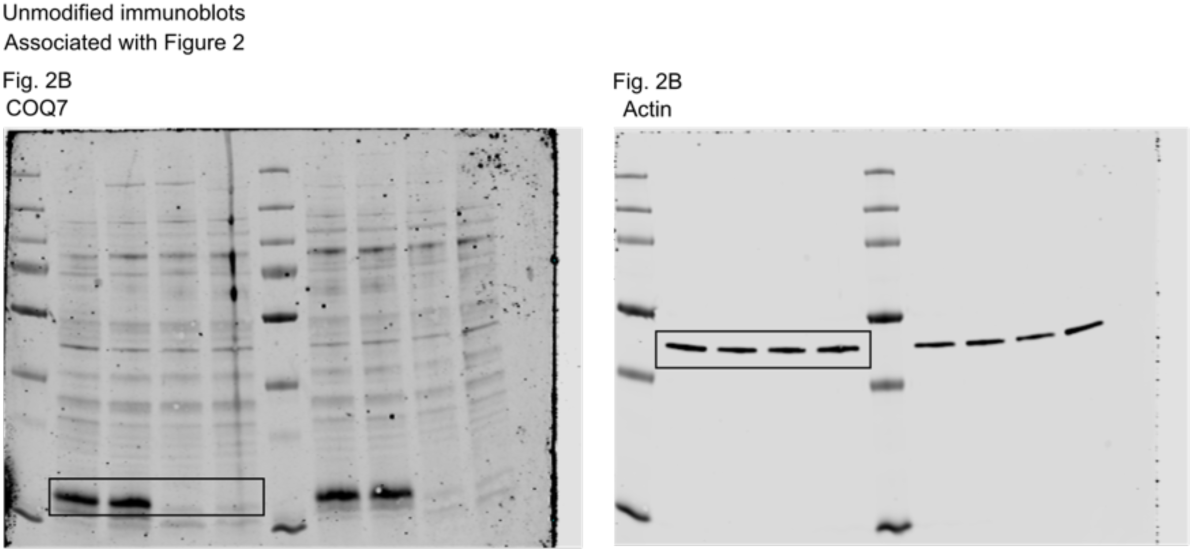
Uncropped immunoblots.

**Source Data Fig. 4:**
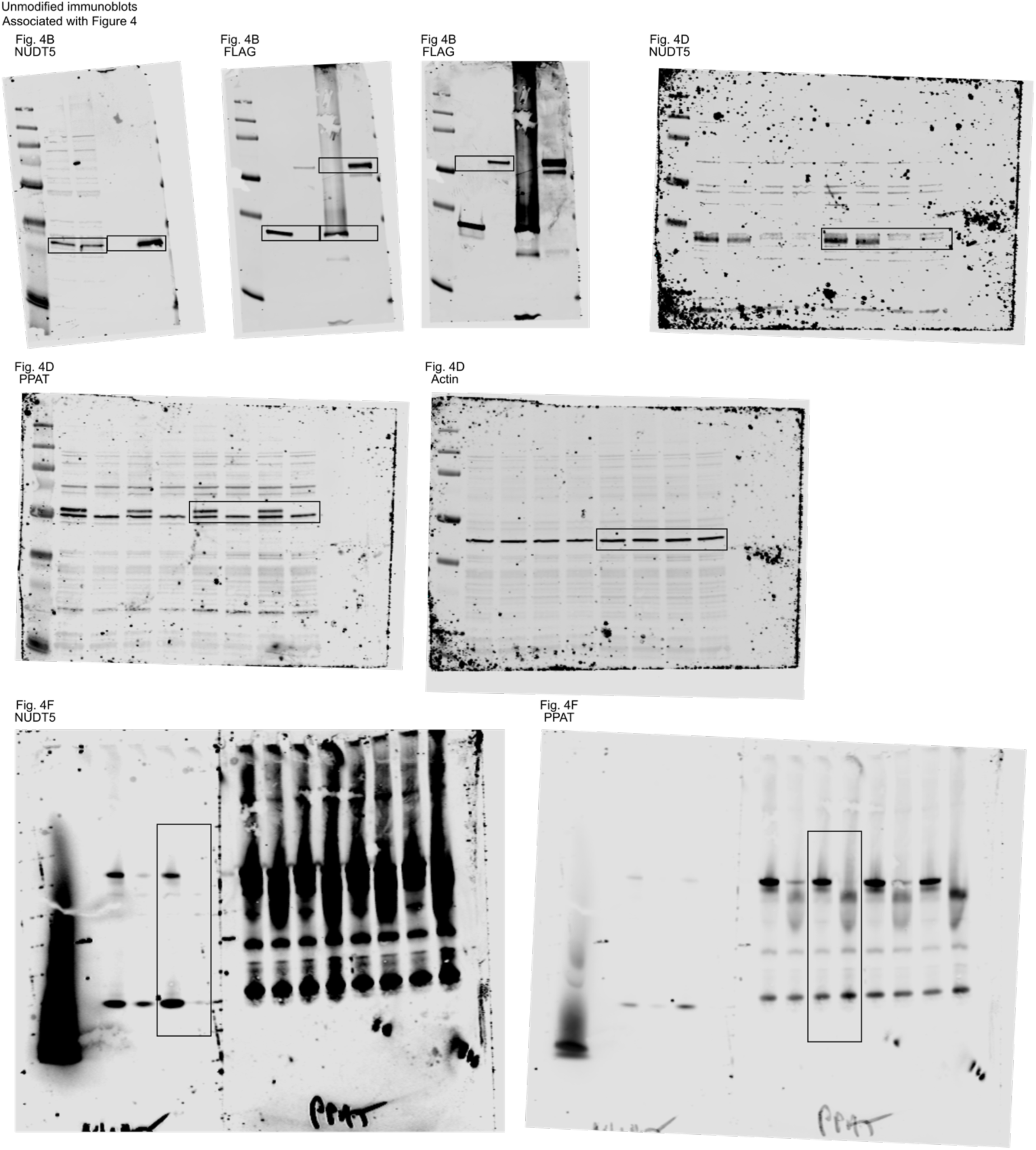
Uncropped immunoblots part 1.

**Source Data Fig. 4:**
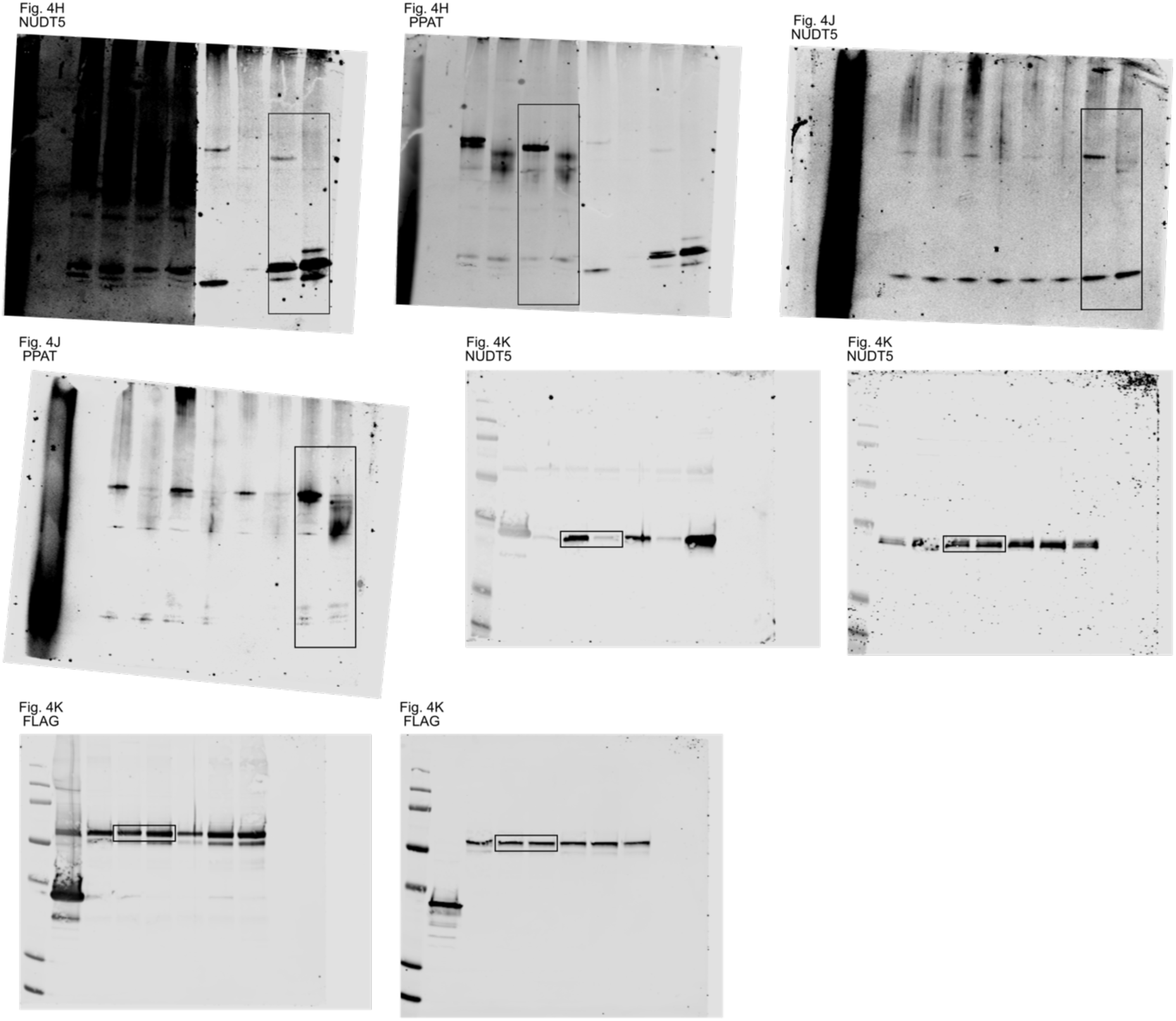
Uncropped immunoblots part 2.

**Source Data Fig. 4:**
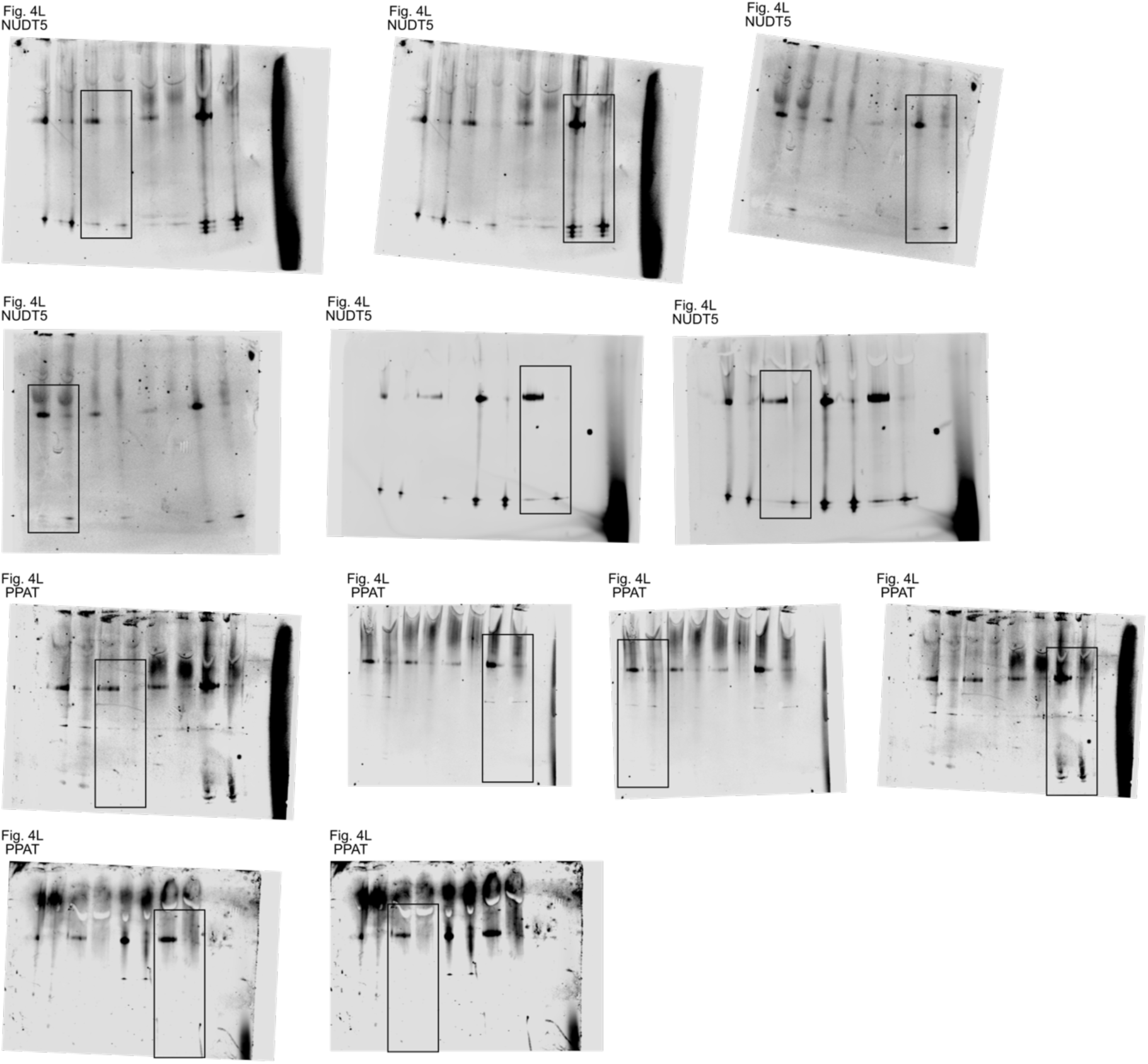
Uncropped immunoblots part 3.

**Source Data Extended Data Fig. 1:**
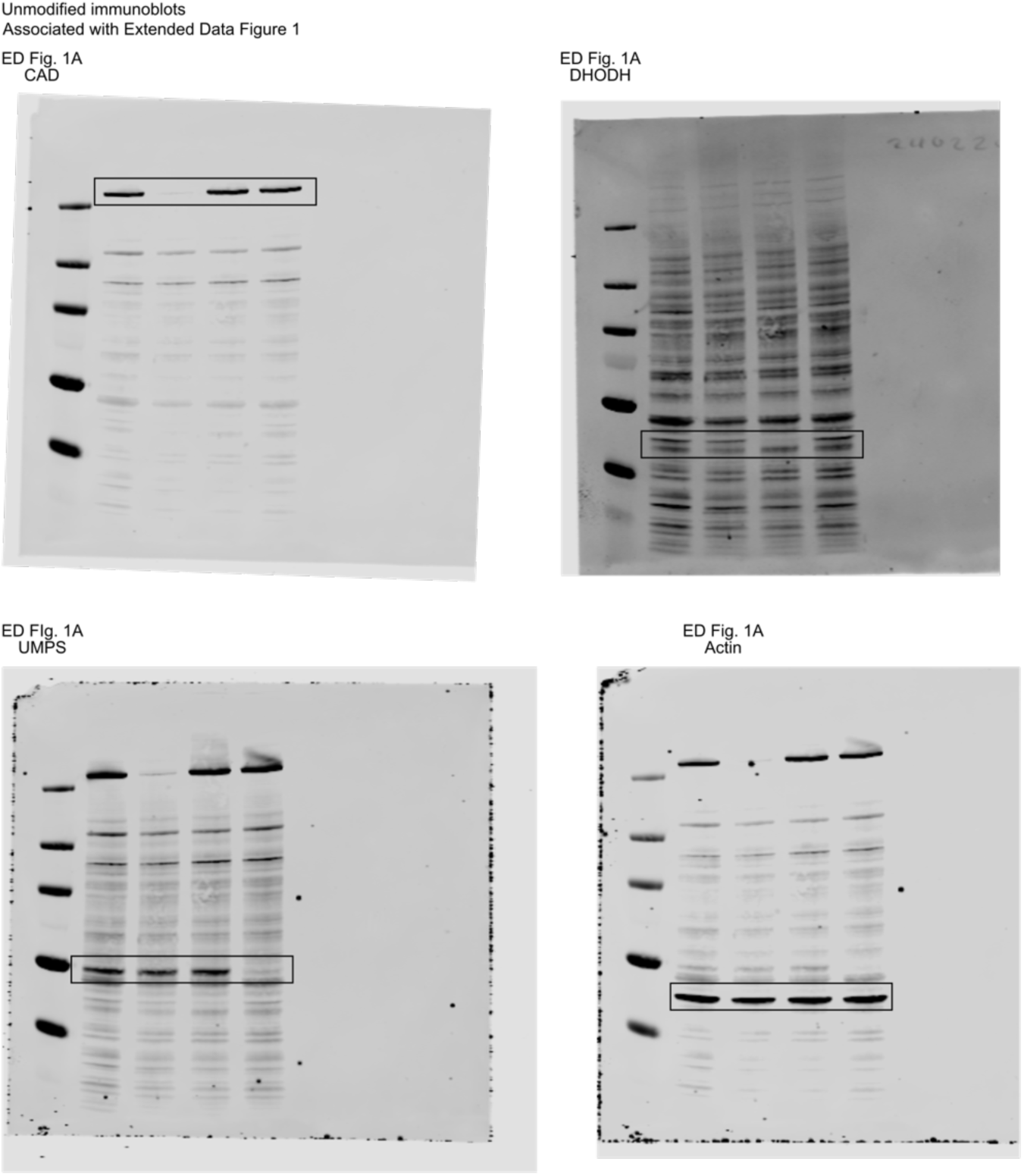
Uncropped immunoblots.

**Source Data Extended Data Fig. 2:**
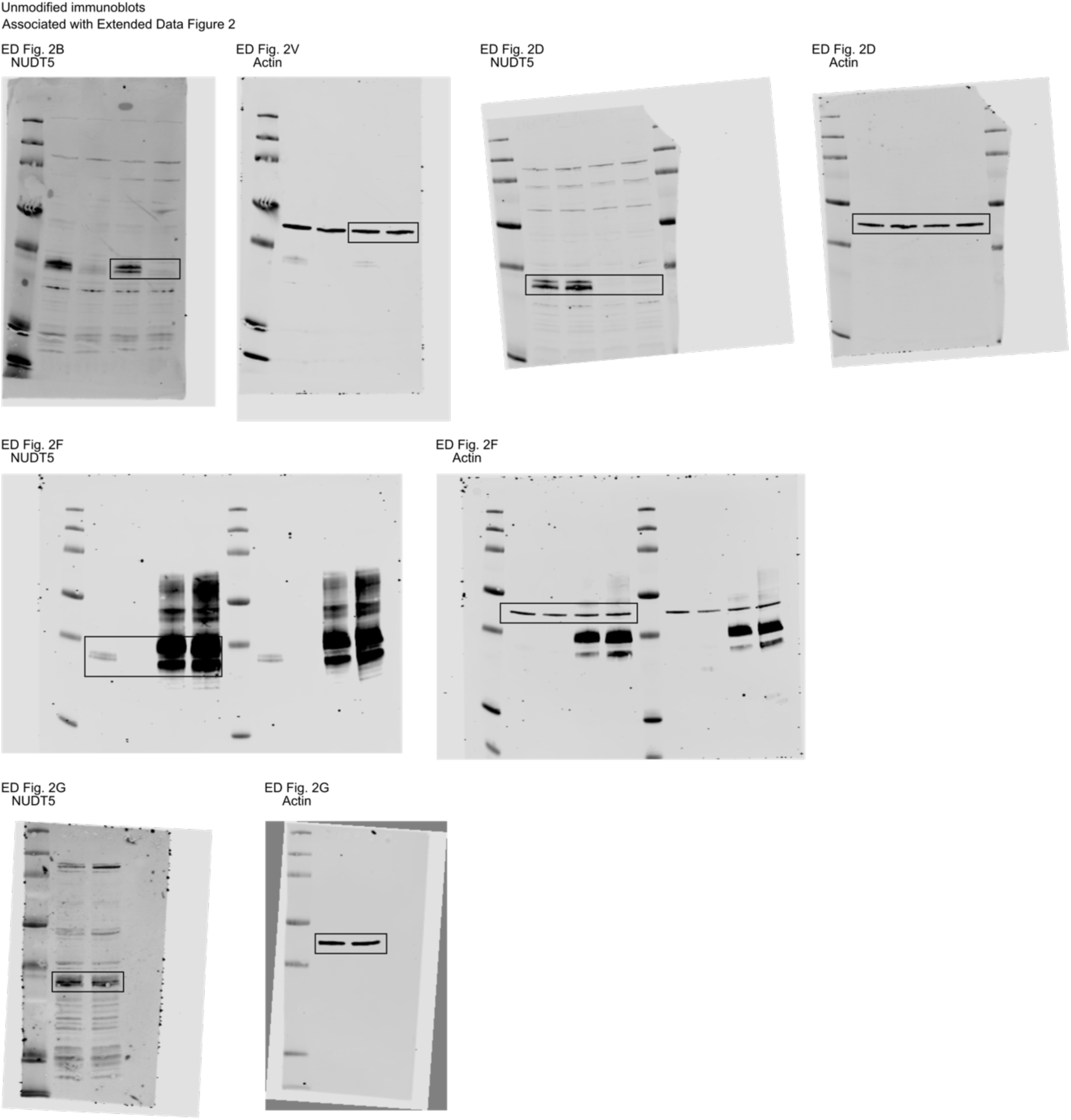
Uncropped immunoblots.

**Source Data Extended Data Fig. 3:**
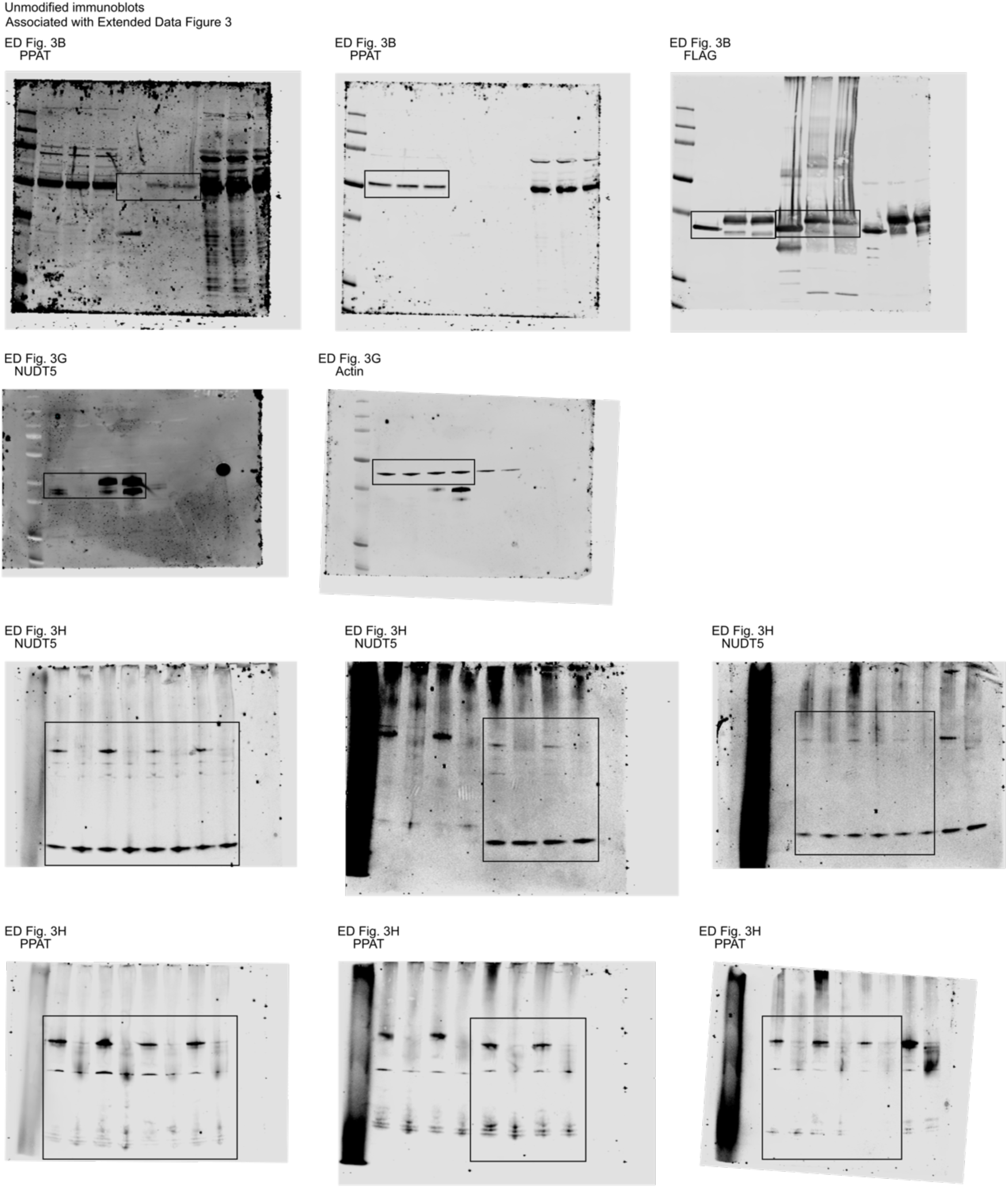
Uncropped immunoblots.

## Materials and Methods

### Animal Experimentation

All animal experiments were approved by the Swiss Cantonal authorities (VD3788) and all relevant ethical regulations were followed. Animals were male C57BL/6J mice aged 12-13 weeks provided with food and water ad libitum and with a standard light–dark cycle of 12 h light exposure. Animals were sacrificed with a high dose of CO_2_, sterilized with 75% ethanol, and collected organs were flash-frozen in liquid nitrogen.

### Cell lines

K562 (ATCC, CCL-243), 293T (ATCC, CRL-3216), MCF7 (ATCC, HTB-22), HeLa (ATCC, CCL-2), and U2OS (ATCC, HTB-96) cells were maintained in DMEM-GlutaMAX (Gibco, 31966021) with 10% fetal bovine serum (FBS, Gibco, A5256701) and 100 U/mL penicillin/streptomycin (BioConcept, 4-01F00-H). UACC-257 (DCTD, CVCL_1779), Jurkat (DSMZ, ACC 282), THP1 (ATCC, TIB-202), U937 (ATCC, CRL-1593.2), and LCL (Coriell, GM12878) cells were maintained in RPMI with 10% FBS (Gibco A5256701) and 100 U/mL penicillin/streptomycin (BioConcept, 4-01F00-H). All cells were cultured under 5% CO_2_ at 37°C. Cells were periodically tested to ensure the absence of mycoplasma.

### Cloning

Gene-specific guide RNAs (sgRNAs) were selected from the two best scoring sequences from the CRISPR-Cas9 screen and cloned into lentiCRISPR v2 vector (Addgene, 52961). The negative control (sgCtrl) was generated using guides targeting *OR2M4* and *OR11A1* that are not expressed in K562. For gene-specific cDNA rescue, sgRNA-resistant gene sequences were cloned into pLV-EF1a-IRES-Puro vector (Addgene, 85132). FLAG-tagged GFP in the same vector was used as a control (Addgene, 201636). The list of sgRNA and cDNA sequences used for cloning can be found in Extended Data Table 4.

### Single cell clones

Individual cloned sgRNA plasmids were electroporated into K562 cells alongside GFP using the Cell Line Nucleofector Kit V (Lonza, VCA-1003) according to the manufacturer’s protocol with T-016 program. Cells were grown for 48 h then stained with Zombie Violet (BioLegend, 423114) and fluorescence-activated single-cell sorting was used to sort GFP^+^ Zombie^−^ cells into flat-bottom 96-well plates at 1 cell per well. Cells were grown for 12 days and wells with single colonies were selected for based on brightfield microscopy. Single cell clones were expanded over 5 weeks and knockouts were verified by immunoblot and by sequencing genomic DNA (gDNA), extracted using the QIAamp DNA kit (Qiagen, 51304) according to the manufacturer’s protocol.

### Virus production and infections

Lentiviruses were produced from 293T cells as previously described^56^. Supernatant was collected 72 h following transfection, filtered through 0.45 µm, and stored at −72°C. For infections, cells at 0.5 × 10^6^ cells/mL with 10 µg/mL polybrene (Sigma-Aldrich, TR-1003) were grown in a 1:1 ratio medium to virus supernatant for 24 h. Selection was performed with 2 µg/mL puromycin (InvivoGen, ant-pr-1) over 48 h. Cells were maintained in standard cell culture medium for 5 days prior to analysis or further experiments. Gene knockout or rescue were confirmed by immunoblot and cDNA rescue with NUDT5 wild-type or E112Q mutant were further verified by sequencing gDNA extracted using the QIAamp DNA kit (Qiagen, 51304), according to the manufacturer’s protocol.

### Growth assays

K562 cells were seeded at 0.05 × 10^6^ cells/mL in black flat-bottom 96-well plates (Thermo Scientific, 137101) for analysis by Prestoblue or in flat-bottom 12-well plates for analysis by cell count. To compare glucose and galactose conditions, test media consisted of DMEM (Gibco, 11966-025) with 1 mM sodium pyruvate (Gibco, 11360070), 2 mM glutamine (BioConcept, 5-10K00-H), 0.2 mM uridine, 10% dialyzed fetal bovine serum (dFBS) (Sigma-Aldrich, F0392), and 100 U/mL penicillin/streptomycin (BioConcept, 4-01F00-H) supplemented with 25 mM glucose or galactose. Base test media for other assays consisted of DMEM (Gibco, 31966021) with 10% dFBS (Sigma-Aldrich, F0392), and 100 U/mL penicillin/streptomycin (BioConcept, 4-01F00-H). This was supplemented with 200 µM uridine, 200 µM cytidine, 25 µM thymidine, volumetric equivalent of water, or a 5-fold dose response curve of chemotherapy drugs in DMSO. Drugs purchased from MedChemExpress were 5-fluorouridine (HY-107856), clofarabine (HY-A0005), fludarabine-phosphate (HY-B0028), and gemcitabine (HY-17026), and from Sigma-Aldrich were 5-fluorouracil (F6627) and 6-thioguanine (A4882). Cells were grown for 5 days under test conditions and proliferation was determined either using the Prestoblue dye (Invitrogen, A13262) and measuring fluorescence (ex/em 560/590 nm) with the BioTek Synergy Plate Reader (Agilent Technologies) after 1.5 h incubation at 37°C, or using trypan-blue based cell counting (Vi-cell Blu counter, Beckman Coulter). Background values were subtracted from Prestoblue data prior to analysis.

### Denaturing polyacrylamide gel electrophoresis (SDS-PAGE)

Cells were harvested, washed in PBS, and lysed by 10 min incubation on ice in RIPA buffer (25 mM Tris pH 7.5, 150 mM NaCl, 0.1% SDS, 0.1% sodium deoxycholate, 1% NP40) with 1:100 protease inhibitor (Thermo Scientific, 87786) and 1:500 nuclease (Thermo Scientific, 88702). Protein concentration was quantified using DC Protein Assay Kit II (Bio-Rad, 5000112). Proteins were separated by SDS-PAGE on Novex Tris-Glycine Mini Protein Gels (Invitrogen, XP08160BOX and XP10200BOX) and were transferred to nitrocellulose membranes using a wet transfer chamber with buffer consisting of 0.302% (w/v) Tris base, 1.44% (w/v) glycine, and 20% ethanol in water. Transfer was verified by Ponceau S Staining Solution (Thermo Scientific, A40000278).

### Immunoblotting

Immunoblotting was performed with 5% milk or 5% bovine serum albumin (Sigma-Aldrich, A9647) in TBS (20 mM Tris-HCl pH 7.4, 150 mM NaCl) with 0.1% tween-20 (TBS-tween) and 1:500-1,000 primary antibodies or 1:5,000 secondary antibodies (LI-COR, 102673-330, 102673-328, or 102673-408). Washes were performed in TBS-tween. Membranes were imaged by fluorescence detection at 700 and 800 nm with Odyssey CLx Imager (LI-COR). For incubation with additional antibodies, membranes were incubated 15 min in mild stripping buffer (15% (w/v) glycine, 1% tween-20, 0.1% SDS, in water pH 2.2), washed in water, and re-blocked. Primary antibodies were actin (Sigma-Aldrich, A3853), CAD (Sigma-Aldrich, HPA057266), COQ7 (Proteintech, 15083-1-AP), DHODH (Sigma-Aldrich, HPA010123), FLAG M2 (Addgene, 194502-rAb), NUDT5 (Sigma-Aldrich, HPA019827), PPAT (Proteintech, 15401-1-AP), and UMPS (Sigma-Aldrich, HPA036178).

### Uridine-sensitized CRISPR-Cas9 screening

Genome-wide CRISPR-Cas9 screening was performed in K562 cells using the Brunello lentiviral library^36^ as previously described^57^. Briefly, K562 cells were infected in duplicate at 500 cells per sgRNA with a multiplicity of infection of 0.3 in the presence of 10 µg/mL polybrene (Sigma-Aldrich, TR-1003). After 24 h, cells were selected with 2 µg/mL puromycin (InvivoGen, ant-pr-1) for 48 h. On day 7 post-infection an aliquot was frozen for comparative analysis. At this time, cells were plated at 10^5^ cells/mL (equivalent to 1,000 cells per sgRNA) in DMEM-GlutaMAX (Gibco, 31966021) with 10% dFBS (Sigma-Aldrich, F0392), 100 U/mL penicillin/streptomycin (BioConcept, 4-01F00-H), and either 200 µM uridine or volumetric equivalent in sterile water. Cells were passaged every 3 days for 3 weeks and 1,000 cells per sgRNA were harvested on day 28 post infection (21 days following media switch). Genomic DNA was extracted using NucleoSpin Blood XL kit (Machery-Nagel, 740954.20), according to the manufacturer’s protocol. Barcode sequencing, mapping, and read count were performed by the Genome Perturbation Platform (Broad Institute).

### Metabolomics analyses

#### Cell preparation

Cells were grown for at least 5 days in DMEM-GlutaMAX (Gibco, 31966021) with 10% dFBS (Sigma-Aldrich, F0392) and 100 U/mL penicillin/streptomycin (BioConcept, 4-01F00-H). PPAT knockout cells were further supplemented with 200 µM inosine. Treatment with 10 µM TH5427 (Tocris Bioscience, 6534) or DMSO was carried out over the last 36 h. Prior to harvest, 10^7^ cells per replicate were transferred to fresh media for 4 h at 37°C. Cells were harvested, washed with PBS, and centrifuged for 1 min at 2,000 g at 4°C. Supernatant was discarded and nitrogen vapor was used to displace ambient air. Pellets were flash-frozen in liquid nitrogen.

#### Targeted metabolomics

Metabolites were extracted using successive freeze-thaw in methanol. Cell pellets were first resuspended in 200 µL of pre-cooled 100% (v/v) methanol containing 1µM of C[13] labelled 4-hydroxybenoic acid as an internal control and then immediately frozen in liquid nitrogen. After brief thawing, samples were centrifuged 5 min at 8,000 g and supernatant was collected. This process was repeated twice on the remaining pellet first with 200 µL 100% (v/v) methanol and then with 100 µL 100% (v/v) water, pooling the supernatants for a final concentration of 80% (v/v) methanol. Pooled supernatants were dried under vacuum, reconstituted in 50 µL of 50% (v/v) acetonitrile:water, and moved into amber glass vials for analysis.

Liquid chromatography coupled to mass spectrometry (LC-MS) analysis was performed using a Thermo Vanquish Horizon UHPLC system coupled to a Thermo Exploris 240 Orbitrap mass spectrometer. For LC separation, a Vanquish binary pump system (Thermo Scientific) was used with a Waters Atlantis Premier BEH Z-HILIC column (100 mm × 2.1 mm, 1.7 µm particle size) held at 35°C under 300 μL/min flow rate. Mobile phase A consisted of 5:95 (v/v) acetonitrile:water with 5 mM ammonium acetate (Sigma Millipore) and 200 µL/L 25% ammonium solution (Sigma Millipore). Mobile phase B consisted of 95:5 (v/v) acetonitrile:water. For each sample run one of two LC methods was used: in the first method, mobile phase B was held at 100% for the first 2 minutes then decreased to 67% over the next 8 minutes. Mobile phase B was then further decreased to 10% and held for 5 minutes. The column was then re-equilibrated for 10 minutes at 100% B before the next injection. In the second method, mobile phase B was held at 100% for the first 3.5 minutes then decreased to 10% over the next 2 minutes and held for 8 minutes. The column was then re-equilibrated for 5 minutes at 100% B before the next injection. 1 µL of sample was injected by a Vanquish Split Sampler HT autosampler (Thermo Scientific) while the autosampler temperature was kept at 4°C. The samples were ionized by a heated electrospray ionization (ESI) source kept at a vaporizer temperature of either 350°C or 200°C depending on the LC method used. Sheath gas was set to 50 units, auxiliary gas to 8 units, sweep gas to 1 unit, and the spray voltage was set to 2,500 V using negative mode. The inlet ion transfer tube temperature was kept at 325°C with 70% RF lens. The identity and retention time of targeted metabolites was first validated using commercial standards when available or unique MS^2^ fragments from a metabolite library (mzcloud.org) when not. Quantification of experimental samples was performed using either parallel reaction monitoring (PRM) with a HCD (30%, 40%, 50%), targeting selected ion fragments generated from the fragmentation of the hydrogen loss (H-) ion or targeted single ion monitoring. Full list of reported precursor and fragment ions is given in Extended Data Table 5. Peak integration was performed using Tracefinder 5.1 (Thermo Scientific).

#### Targeted lipidomics

Frozen cell pellets were thawed on ice, then 150 mM KCl (50 µL) was added to each sample, followed by ice-cold methanol (600 µL) with 1 µM CoQ_8_ as internal standard (Avanti Polar Lipids). The samples were vortexed 10 min at 4°C to lyse the cells. Ice-cold petroleum ether (400 µL) was added to extract the lipids, and the samples were vortexed again for 3 min at 4°C. Samples were centrifuged 3 min at 1,000 g at 21°C and the top petroleum ether layer was collected in a new tube. The petroleum ether extraction was repeated a second time, with the petroleum ether layer from the second extraction combined with that from the first. The extracted lipids were dried under argon before being resuspended in isopropanol (40 µL) and transferred to an amber glass vial (Supelco, QSertVial, 12 × 32 mm, 0.3 mL).

LC-MS analysis was performed using a Thermo Vanquish Horizon UHPLC system coupled to a Thermo Exploris 240 Orbitrap mass spectrometer. For LC separation, a Vanquish binary pump system (Thermo Scientific) was used with a Waters Acquity CSH C18 column (100 mm × 2.1 mm, 1.7 µm particle size) held at 35°C under 300 μL/min flow rate. Mobile phase A consisted of 5 mM ammonium acetate in 70:30 (v/v) acetonitrile:water with 125 μL/L acetic acid. Mobile phase B consisted of 5 mM ammonium acetate in 90:10 (v/v) isopropanol:acetonitrile with the same additive. For each sample run, mobile phase B was initially held at 2% for 2 min and then increased to 30% over 3 min. Mobile phase B was further increased to 50% over 1 min and 85% over 14 min and then raised to 99% over 1 min and held for 4 min. The column was re-equilibrated for 5 min at 2% B before the next injection. Five microliters of the sample were injected by a Vanquish Split Sampler HT autosampler (Thermo Scientific), while the autosampler temperature was kept at 4°C. The samples were ionized by a heated ESI source kept at a vaporizer temperature of 350°C. Sheath gas was set to 50 units, auxiliary gas to 8 units, sweep gas to 1 unit, and the spray voltage was set to 3,500 V for positive mode and 2,500 V for negative mode. The inlet ion transfer tube temperature was kept at 325°C with 70% RF lens. For targeted analysis, the MS was operated in parallel reaction monitoring mode with polarity switching acquiring scheduled, targeted scans to CoQ_10_ H^+^ adduct (m/z 863.6912), CoQ_10_ NH4^+^ adduct (m/z 880.7177), CoQ_8_ H^+^ adduct (m/z 727.566), CoQ_8_ NH4^+^ adduct (m/z 744.5935) and CoQ intermediates: DMQ_10_ H^+^ adduct (m/z 833.6806), DMQ_10_ NH4^+^ adduct (m/z 850.7072), and PPHB_10_ H^−^ adduct (m/z 817.6504). MS acquisition parameters include resolution of 45,000, HCD collision energy (45% for positive mode and 60% for negative mode), and 3s dynamic exclusion. Automatic gain control targets were set to standard mode. The resulting CoQ intermediate data were processed using TraceFinder 5.1 (Thermo Scientific). Raw intensity values were normalized to the CoQ_8_ internal standard.

#### Multiple-pathways targeted metabolomics

Cell pellets were extracted with 80% (v/v) methanol, sonicated, and homogenized with ceramic beads (Precellys Cryolys). Lysates were centrifuged 15 min at 15,000 g at 4°C and the supernatant was evaporated to dryness. Dried extracts were reconstituted in methanol according to total protein content as measured by BCA assay. Samples were analyzed by ultra-high performance liquid chromatography coupled to tandem mass spectrometry (UHPLC-MS/MS), using the Triple Quadrupole mass spectrometer (6495 iFunnel, Agilent Technologies) and dynamic Multiple Reaction Monitoring (dMRM) acquisition mode, following previously described methods^58,59^. Two complementary liquid chromatography modes coupled to positive and negative electrospray ionization MS, respectively, were used to maximize the metabolome coverage^60^.

Raw UHPLC-MS/MS data were processed using the MassHunter Quantitative Analysis software vB.07.00 (Agilent Technologies). Extracted ion chromatogram areas for MRM transitions were used for relative quantification. The data, consisting of peak areas of detected metabolites across all samples, were processed and filtered depending on the coefficient of variation (CV) evaluated across quality control samples that were analyzed periodically throughout the batch. Peaks with analytical variability above CV of 30% were discarded.

### Native polyacrylamide gel electrophoresis

For cells: Cells were harvested, washed with PBS, and lysed by 10 min incubation on ice in 1X Native Buffer (NativePAGE Sample Prep kit, Invitrogen, BN2008), 1% digitonin (Invitrogen, BN2006), 1:100 protease inhibitor (Thermo Scientific, 87786), 1:500 nuclease (Thermo Scientific, 88702), and 10 mM individual metabolites as indicated.

For mouse tissues: Frozen mouse tissue was homogenized (gentleMACS Octo Dissociator, Miltenyi Biotec) in 50 mM Tris-HCl pH 7.4, 150 mM NaCl, 1 mM MgCl_2_, 1% NP-40, 1:100 protease inhibitor (Thermo Scientific, 87786), and 1:500 nuclease (Thermo Scientific, 88702) then centrifuged 10 min at 3,000 g. Supernatant was diluted in the same buffer and 10 mM metabolites were added as indicated.

Samples with metabolite supplementation were incubated 1-2 h at 4°C with gentle agitation. Lysates were centrifuged 10 min at 20,000 g at 4°C and the supernatant was saved. Protein concentration was quantified using DC Protein Assay Kit II (Bio-Rad, 5000112). Proteins were separated on a 4-16% native gel (Invitrogen, BN1004BOX) with 1X anode buffer (NativePAGE Running Buffer kit, Invitrogen, BN2007) and transferred to nitrocellulose membranes using a wet transfer chamber and buffer consisting of 0.302% (w/v) Tris base, 1.44% (w/v) glycine, 20% ethanol, and 0.05% (w/v) SDS in water. Transfer was verified by Ponceau S Staining Solution (Thermo Scientific, A40000278). Membranes were incubated 5 min in 8% acetic acid and washed with water before proceeding to immunoblotting.

### Oxygen consumption rate

K562 cells were seeded at 125’000 cells/well in Seahorse XF DMEM (Agilent Technologies, 103575-100) supplemented with 25 mM glucose and 2 mM glutamine, centrifuged 1 min at 100 g, and incubated 1 h at 37°C. Oxygen consumption rate was measured by the Agilent Seahorse XFe96 Extracellular Flux Analyzer (Agilent Technologies) using the program XF Cell MitoStress Test and successive treatment with 2 µM oligomycin, 1.5 µM CCCP, and 1.6 µM antimycin. Data were analyzed using Seahorse Wave Desktop Software (Agilent Technologies).

### Immunoprecipitation

293T cells were transfected with FLAG-tagged NUDT5, NUDT5 mutants, PPAT, or GFP as control using lipofectamine 2000 (Invitrogen, 11668019) according to the manufacturer’s protocol. Transfected cells were grown to 80% confluency, harvested by scraping, washed with PBS, and lysed by 15-30 min incubation on ice in IP-buffer (50 mM Tris-HCl pH 7.4, 150 mM NaCl, 1 mM MgCl_2_, 1% NP-40) with 1:100 protease inhibitor (Thermo Scientific, 87786) and 10 mM metabolites as indicated. Lysates were spun 10 min at 20,000 g at 4°C and supernatants were transferred to new tubes. 1% volume was kept aside for inputs. Anti-FLAG M2 magnetic beads (Millipore, M8823) were washed in IP-buffer and incubated with samples overnight at 4°C with gentle agitation. Beads were washed five times in IP-buffer and changed twice to new tubes. 0.1 mg/mL 3xFLAG peptide (Sigma-Aldrich, F4799) in TBS was added to beads in two steps and each time incubated 30 min at 4°C with gentle agitation, supernatant was saved on ice. 100 µL trichloroacetic acid (Sigma-Aldrich, T9159) was added to the supernatant and incubated 30 min at 4°C. Samples were centrifuged 20 min at 20,000 g at 4°C, supernatant was discarded, and pellets were washed in −20°C acetone. Samples were centrifuged 10 min at 20,000 g, supernatant was discarded, and pellets were heated at 55°C until dry. Pellets were resuspended in 2X SDS sample buffer for analysis by SDS-PAGE or mass spectrometry.

### Proteomics analyses

#### Global proteomics

Cells were cultured 5 days in DMEM-GlutaMAX (Gibco, 31966021) with 10% dFBS (Sigma-Aldrich, F0392) and 100 U/mL penicillin/streptomycin (BioConcept, 4-01F00-H). 5 × 10^6^ cells were harvested, washed with PBS, and pellets were flash-frozen in liquid nitrogen. Protein extraction, library construction and analyses, and sample analyses were all carried out as described previously^61^, with minor changes. Briefly, proteins were extracted using a modified iST method and dried by centrifugal evaporation. 1/8 samples were pooled for library construction and were fractionated by off-line basic reversed-phase fractionation (bRP). Dried bRP fractions were redissolved in 30 µL 2% acetonitrile with 0.5% TFA and 6 µL were injected for LC-MS/MS analysis. LC-MS/MS analyses were carried out on a TIMS-TOF Pro mass spectrometer (Bruker) interfaced through a nanospray ion source to an Ultimate 3000 RSLCnano HPLC system (Dionex) using data-dependent acquisition (DDA) for library construction and data-independent acquisition (DIA) for sample analysis.

Raw Bruker MS data were processed directly with Spectronaut 17.5 (Biognosys). A library was constructed from the DDA bRP fraction data using the annotated SWISSPROT human proteome database of 2022-01-07, containing 20,375 sequences, using parameters described elsewhere^61^. The library created contained 119,959 precursors mapping to 86,501 stripped sequences, of which 82,494 were proteotypic. These corresponded to 7,853 protein groups or 7,954 proteins. Of these, 830 were single hits (one peptide precursor). In total, 708,314 fragments were used for quantitation. Peptide-centric analysis of DIA data was done with Spectronaut 17.5 using the library described above and criteria described elsewhere^61^. Overall, 112,958 precursors were quantified in the dataset, mapped to 7,262 protein groups or 7,343 proteins. 99,738 precursors (7,076 protein groups) had full profiles, i.e. were quantified in all samples. The average number of data points was 7.2.

#### IP-MS proteomics

Samples were immunoprecipitated as described above. Digestion was carried out using the SP3 method^62^ with magnetic Sera-Mag Speedbeads (Cytiva, 45152105050250). Proteins were alkylated with 32 mM idoacetamine for 45 min at 21°C in the dark. Beads were added at a ratio 10:1 (w/w) to samples and proteins were precipitated on beads with ethanol at a final concentration of 60%. After three washes with 80% ethanol, beads were digested in 50 µL of 100 mM ammonium bicarbonate with 1 µg trypsin (Promega V5073) and incubated 1 h at 37°C. The same amount of trypsin was added to samples for an additional 1 h incubation. Supernatant was then recovered, transferred to new tubes, acidified with formic acid at 0.5% final concentration, and dried by centrifugal evaporation. In order to remove traces of SDS, two sample volumes of isopropanol containing 1% TFA were added to the digests, and samples were desalted on a strong cation exchange plate (Oasis MCX, Waters) by centrifugation. Digests were washed with 1% TFA in isopropanol then 0.1% formic acid with 2% acetonitrile. Peptides were eluted in 200 µL of 80% MeCN, 19% water, 1% (v/v) ammonia, and dried by centrifugal evaporation.

LC-MS/MS analyses were carried out on a TIMS-TOF Pro mass spectrometer (Bruker) interfaced through a nanospray ion source to an EvoSep One liquid chromatography system (EvoSep). Peptides were separated on a reversed-phase 15 cm C18 column (150 µm ID, 1.5 µm, EvoSep EV1137) at a flow rate of 0.22 µL/min with a 15 sample per day method (runtime 88 min, solvents were water and acetonitrile with 0.1% formic acid, useful gradient 0-35%). DDA was carried out using a method similar to the standard TIMS PASEF method^63^ with ion accumulation for 100 ms for each survey MS1 scan and TIMS-coupled MS2 scans. Duty cycle was kept at 100%. Precursor ions were chosen within the ion mobility range from 1/k0 = 0.8 and between m/z 400-1200. Up to ten precursors were targeted per TIMS scan. Precursor isolation was done with a 2 or 3 m/z window below or above m/z 800, respectively. The minimum threshold intensity for precursor selection was 2500. If the inclusion list allowed it, precursors were targeted more than once to reach a minimum target intensity of 20,000. Collision energy was ramped linearly based uniquely on the 1/k0 values from 20 (at 1/k0 = 0.6) to 59 eV (at 1/k0 = 1.6). Total duration of a scan cycle including one survey and 10 MS2 TIMS scans was 1.16 s. Precursors could be targeted again in subsequent cycles if their signal increased by a factor of 4.0 or more. After selection in one cycle, precursors were excluded from further selection for 60 s. Mass resolution in all MS measurements was approximately 35,000.

Data were analyzed with MaxQuant v2.4.11.0^64^ incorporating the Andromeda search engine^65^. Cysteine carbamidomethylation was selected as fixed modification while methionine oxidation and protein N-terminal acetylation were specified as variable modifications. The sequence databases used for searching were the SWISSPROT human proteome database of 2024-02-14, containing 82,499 sequences, and a “contaminant” database containing the most usual environmental contaminants and enzymes used for digestion. Mass tolerance was 4.5 ppm on precursors (after recalibration) and 20 ppm on MS/MS fragments. Both peptide and protein identifications were filtered at 1% FDR relative to hits against a decoy database built by reversing protein sequences.

### ADP/ATP assay

Cells were harvested, washed in PBS, and seeded at 10^6^ cells/mL in white flat-bottom 96-well plates (Thermo Scientific, 136101). Cellular ADP and ATP levels were measured using the ADP/ATP Ratio Assay Kit (Sigma-Aldrich, MAK135) according to the manufacturer’s protocol with the BioTek Plate Reader (Agilent Technologies).

### (Hypo)xanthine assay

Cells were harvested, washed with PBS, and lysed by 10 min incubation on ice in Assay Buffer (Abcam, ab155900) at 2 × 10^7^ cells/mL. (Hypo)xanthine concentration was measured using the fluorometric protocol of the Xanthine/Hypoxanthine Assay Kit (Abcam, ab155900) in black flat-bottom 96-well plates (Thermo Scientific, 137101). Fluorescence was measured at ex/em 535/587 nm with the BioTek Synergy Plate Reader (Agilent Technologies).

### Data analysis

#### CRISPR screen data analysis

For analysis by Z-score, sgRNA read count data were processed using an approach previously described^22^. Briefly, data were normalized to reads per million and transformed to Log_2_ space. A Log_2_ fold-change for each sgRNA in each media condition was determined relative to the mean day 7 control read count (prior to media switch). The mean Log_2_ fold-change was calculated across sgRNAs for each gene, and results were averaged across two infection replicates. Low-expression genes (TPM < 1 in DepMap 24Q2 dataset^66^) were used to define the mean and standard deviation of a null distribution for each media condition and Z-scores for each gene were defined based on this distribution. These low-expression genes and their corresponding sgRNA were excluded from further analyses. sgRNA read count data from day 28 (day 21 post infection) were used as input for MaGeCK v0.5.9.2^23^ with default parameters and using the media condition supplemented with uridine as the reference.

#### Gene Set Enrichment Analysis

Gene set enrichment analysis was performed using GSEA^24,25^ v4.2.2 on genes ranked according to ΔZ = Z_-uridine_ – Z_+uridine_ or by gene Log_2_ fold-change as calculated by MaGeCK. Enrichment was performed against the GO Biological Processes database c5.go.bp.v2023.2.Hs.symbols without collapse. Maximum size exclusion was set to 50 and minimum to 2, all other parameters were kept as default. The top 50 negatively enriched pathways were manually annotated for their relationship to either pyrimidine biosynthesis or CoQ metabolism.

#### Metabolite Set Enrichment Analysis

Metabolites were mapped to their KEGG^67^ IDs using MetaboAnalyst v6.0 “Compound ID Conversion” tool (https://www.metaboanalyst.ca/). Metabolites that were not successfully found were mapped manually where possible and were otherwise excluded. KEGG pathway identifiers for Metabolism pathways were taken from their website (https://www.genome.jp/kegg/pathway.html on 2024-10-24), excluding the subsections “Xenobiotics degradation and metabolism” and “Chemical structure transformation maps”. The list of metabolites for each pathway was obtained using the KEGGREST v1.44.1 package (https://bioconductor.org/packages/KEGGREST) in R v4.4.1. These data were used to generate a metabolite database and four additional pathways were added manually, delimiting mature purines or pyrimidines from the intermediate metabolites of their respective *de novo* synthesis pathways. Metabolite set enrichment analysis was performed using GSEA^24,25^ v4.2.2 on metabolites ranked according to their Log_2_ fold-change (sgNUDT5 / sgCtrl) in abundance. Enrichment was performed against the custom database described above without collapse. Maximum size exclusion was set to 500 and minimum to 3, all other parameters were kept as default.

#### Global proteomics

Analyses were done with the Perseus software package v1.6.15.0^68^. Contaminant proteins were removed, data were Log_2_ transformed, and only proteins quantified in at least four samples of one group were kept. Missing values were imputed based on normal distribution using Perseus default parameters. Student t-tests were carried out and the Log_2_ fold-change over control samples (sgCtrl) was calculated.

#### IP-MS proteomics

Analyses were done using an in-house developed software tool (available on https://github.com/UNIL-PAF/taram-backend). Contaminant proteins were removed, data were Log_2_ transformed, and only proteins quantified in at least two samples of one group were kept, resulting in 2,949 protein groups. Missing values were imputed based on a normal distribution with a width of 0.3 standard deviations, down-shifted by 1.8 standard deviations relative to the mean. Student t-tests were carried out and the Log_2_ fold-change over control samples (GFP-3xFLAG) was calculated.

#### Other statistical analyses

Statistical analyses as described in the figure legends were performed using Prism 10 (GraphPad Software) and exact p-values are shown where p < 0.05.

